# Assessing the impact of grassland management on landscape multifunctionality

**DOI:** 10.1101/2020.07.17.208199

**Authors:** M. Neyret, M. Fischer, E. Allan, N. Hölzel, V.H. Klaus, T. Kleinebecker, J. Krauss, G. Le Provost, S. Peter, N. Schenk, N.K. Simons, F. van der Plas, J. Binkenstein, C. Börschig, K. Jung, D. Prati, D. Schäfer, M. Schäfer, I. Schöning, M. Schrumpf, M. Tschapka, C. Westphal, P. Manning

## Abstract

Land-use intensification has contrasting effects on different ecosystem services, often leading to land-use conflicts. While multiple studies have demonstrated how landscape-scale strategies can minimise the trade-off between agricultural production and biodiversity conservation, little is known about which land-use strategies maximise the landscape-level supply of multiple ecosystem services (landscape multifunctionality), a common goal of stakeholder communities.

We combine comprehensive data collected from 150 German grassland sites with a simulation approach to identify landscape compositions, with differing proportions of low-, medium-, and high-intensity grasslands, that minimise trade-offs between the six main grassland ecosystem services prioritised by local stakeholders: biodiversity conservation, aesthetic value, productivity, carbon storage, foraging, and regional identity. Results are made accessible through an online tool that provides information on which compositions best meet any combination of user-defined priorities (https://neyret.shinyapps.io/landscape_composition_for_multifunctionality/).

Results show that an optimal landscape composition can be identified for any pattern of ecosystem service priorities. However, multifunctionality was similar and low for all landscape compositions in cases where there are strong trade-offs between services (e.g. aesthetic value and fodder production), where many services were prioritised, and where drivers other than land use played an important role. We also found that if moderate service levels are deemed acceptable, then strategies in which both high and low intensity grasslands are present can deliver landscape multifunctionality. The tool presented can aid informed decision-making by predicting the impact of future changes in landscape composition, and by allowing for the relative roles of stakeholder priorities and biophysical trade-offs to be understood by scientists and practitioners alike.

**Highlights:**

- An online tool identifies optimal landscape compositions for desired ecosystem services
- When the desired services are synergic, the optimum is their common best landscape composition
- When the desired services trade-off, a mix of grassland intensity is most multifunctional
- Such tools could support decision-making processes and aid conflict resolution

**Graphical abstract:** 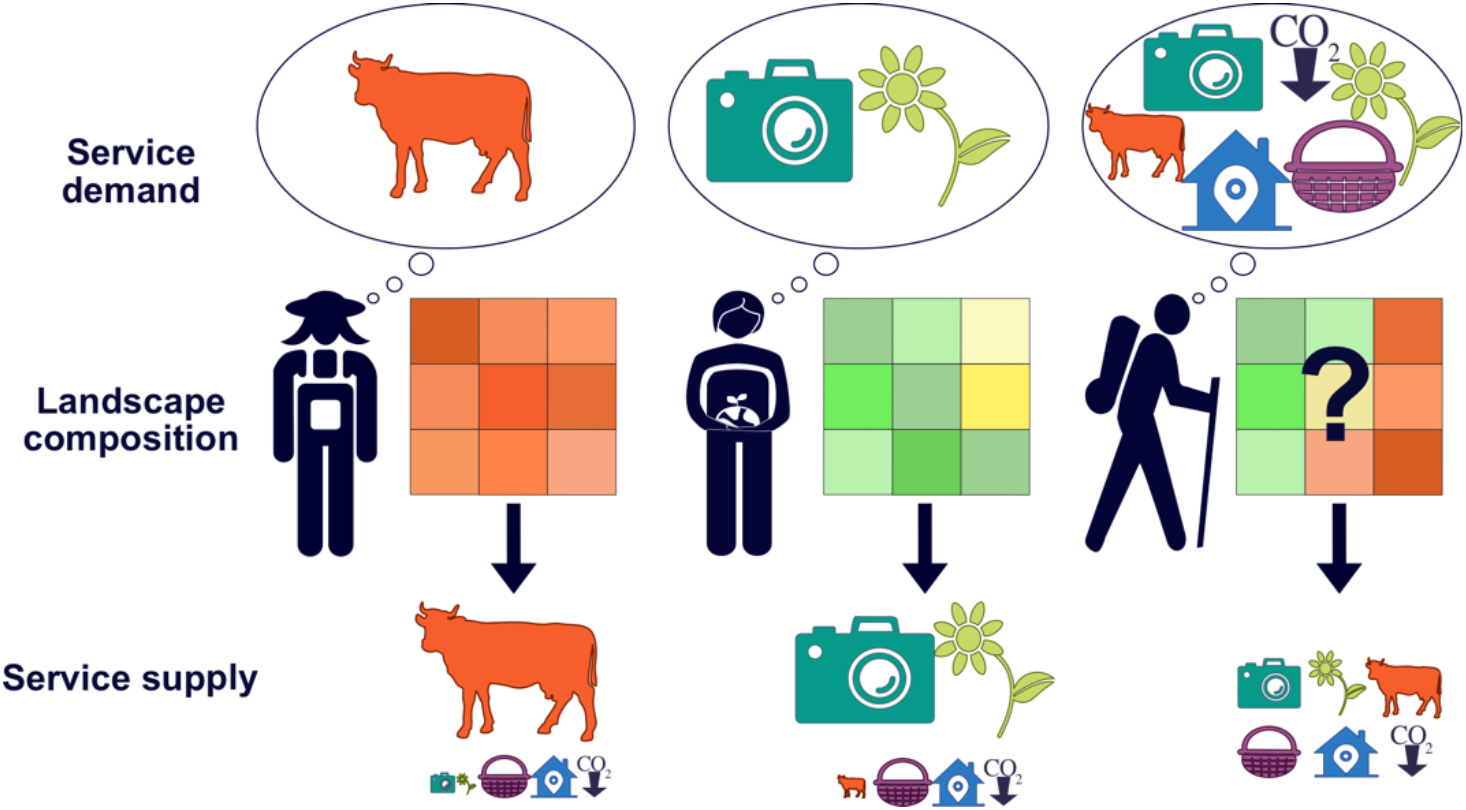

## 1. Introduction

Habitat conversion and land-use intensification are driving biodiversity loss and changes to ecosystem service supply across the world (IPBES 2019). While high land-use intensification promotes a small number of ecosystem services related to food production, it is often detrimental to biodiversity conservation (Bennett et al., 2009; Lavorel et al., 2011; Raudsepp-Hearne et al., 2010) and other regulating or cultural ecosystem services that depend on biodiversity for their delivery (Allan et al., 2015; Clec’h et al., 2019). Such contrasting responses of different ecosystem services to ecosystem drivers often make it impossible to achieve high levels of all desired services at a local and landscape scale (van der Plas et al., 2019). This has led to land-use conflicts, which are becoming increasingly common across the globe (Goldstein et al., 2012).

To date, much of the work on minimising trade-offs between ecosystem services within landscapes has compared a ‘land sparing’ strategy, in which semi-natural high-biodiversity areas and intensive farmland are spatially segregated, and a ‘land sharing’ strategy in which biodiversity conservation and commodity production are co-delivered in a landscape of intermediate intensity, and where these different land uses form a mosaic (Green, 2005). Within this field, most studies have found that land sparing is the best way to achieve high levels of both biodiversity conservation and commodity production (Phalan et al., 2011; Simons & Weisser, 2017). However, multiple studies have also stressed the limitations of the land sharing versus land sparing concept. The framework focuses on just two extreme strategies, and on only two services - commodity production and biodiversity conservation (Bennett, 2017; Fischer et al., 2014), while in reality, most landscapes are expected to provide multiple services, even within a single ecosystem type. This is the case for semi-natural grasslands (*sensu* Bullock et al. 2011), which supply a wide range of highly prioritised ecosystem services including water provision, climate regulation (carbon storage) and recreation services, in addition to food production and biodiversity conservation (Bengtsson et al., 2019). Accounting for these additional ecosystem services could significantly affect which land-use strategy is considered optimal, meaning the best strategy for achieving high levels of multiple services within grassland landscapes remains unknown.

One way of measuring how land use affects the overall supply of multiple services is the ecosystem service multifunctionality approach (Manning et al., 2018). Ecosystem service multifunctionality is defined as the simultaneous supply of multiple prioritised ecosystem services, relative to their human demand (Linders et al. 2021). It builds upon the metrics used in biodiversity-ecosystem functioning research (Allan et al., 2015; Barnes et al., 2017; Byrnes et al 2014) by combining ecological and biophysical data describing the supply of multiple ecosystem services with social data that quantifies the relative priority given by stakeholder groups to each service. The approach also advances on existing methods such as the identification of ecosystem service bundles (Frei et al., 2018; Raymond et al., 2009) by measuring the overall supply of ecosystem services relative to their demand. The resulting multifunctionality metrics can therefore be seen as summarising the overall benefit provided by a system to stakeholders.

Here, we combine the multifunctionality approach with data simulation methods to identify the optimal landscape composition for multiple ecosystem services. This approach involves varying the proportion of land under different intensities in data simulations and measuring the outcome on ecosystem service multifunctionality. We also investigate how the relative priority of services to land users affects the optimal strategy. The analysis was achieved by using ecosystem service data collected at 150 grassland sites that vary in their intensity, found in the three regions of the large-scale and long-term Biodiversity Exploratories project, in Germany. This was utilised in simulations in which artificial ‘landscapes’ of varying composition, in terms of land-use intensity, were assembled from site-level data (Fig. 1). We base our metrics of multifunctionality on six services which are directly linked to final benefits (*sensu* the cascade model, Mace et al., 2012; Fisher & Turner, 2008): fodder production, biodiversity conservation, climate change mitigation, aesthetic value, foraging opportunities and regional identity, covering all the services provided by grasslands that were demanded by the main stakeholder groups, as identified in a social survey (Supplementary Fig. A1). We hypothesised that heterogeneous landscapes composed of both high- and low-intensity sites would have the highest multifunctionality (van der Plas et al., 2019).

**Figure 1.**
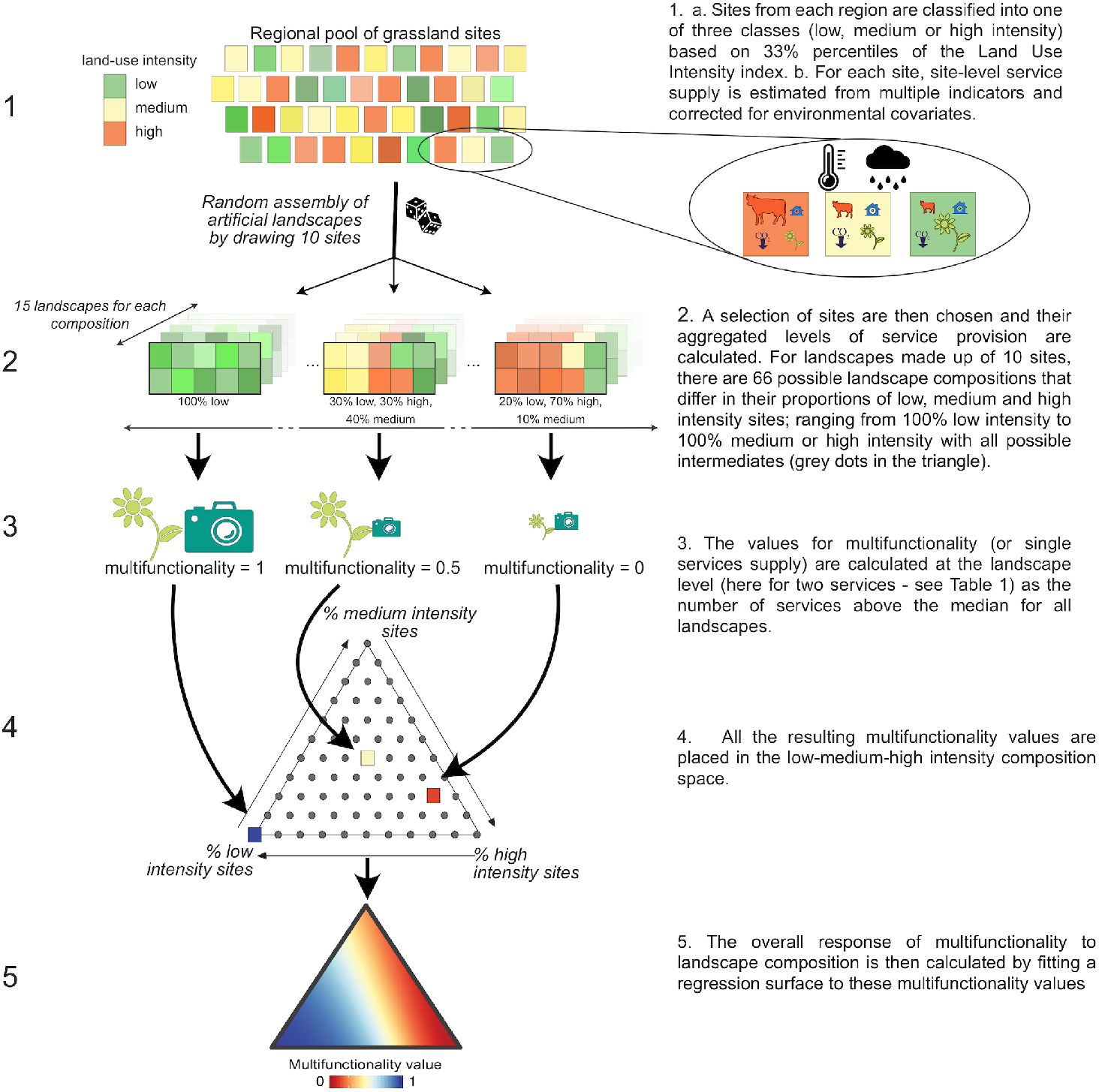
Steps of the analysis.

## 2. Material and methods

### 2.1 Study design

We used data from 150 grassland plots (hereafter sites) studied within the large-scale and long-term Biodiversity Exploratories project in Germany (https://www.biodiversity-exploratories.de/). The sites were located in three regions: the UNESCO Biosphere Area Schwäbische Alb (South-West region), in and around the National Park Hainich (Central region; both are hilly regions with calcareous bedrock), and the UNESCO Biosphere Reserve Schorfheide-Chorin (North of Germany: flat, with a mixture of sandy and organic soils, see Fischer et al. (2010) for details). The regions were selected to be broadly representative of the three main landscape types of Southern, Central and Northern Germany in terms of environmental conditions and management regimes. Sites measured 50 × 50m and were selected to be representative of the whole field they were in. Field size ranged from 3 to 148 ha. The 50 sites of each region spanned the full range of land-use intensity within the region, while minimising variation in potentially confounding environmental factors.

It has been shown that the shape of the response of ecosystem services to their drivers can affect which landscape strategies are identified as optimal (e.g. Phalan et al. 2011), and in our study, the response of each service to land-use intensity differs strongly across regions, even after correcting for the effects of environmental covariates. For instance, biodiversity in the North region is inherently and uniformly low, hence an almost ‘flat’ biodiversity response to LUI is observed, while diversity declines strongly between low and high intensity in the other two regions (see Results, Fig. 2a). As a result, the optimal strategy is likely to be region-specific and identifying optimal strategies based on ‘averaged’ responses could give dangerously misleading results. Therefore, we chose to run all analyses at the regional level.

**Figure 2.**
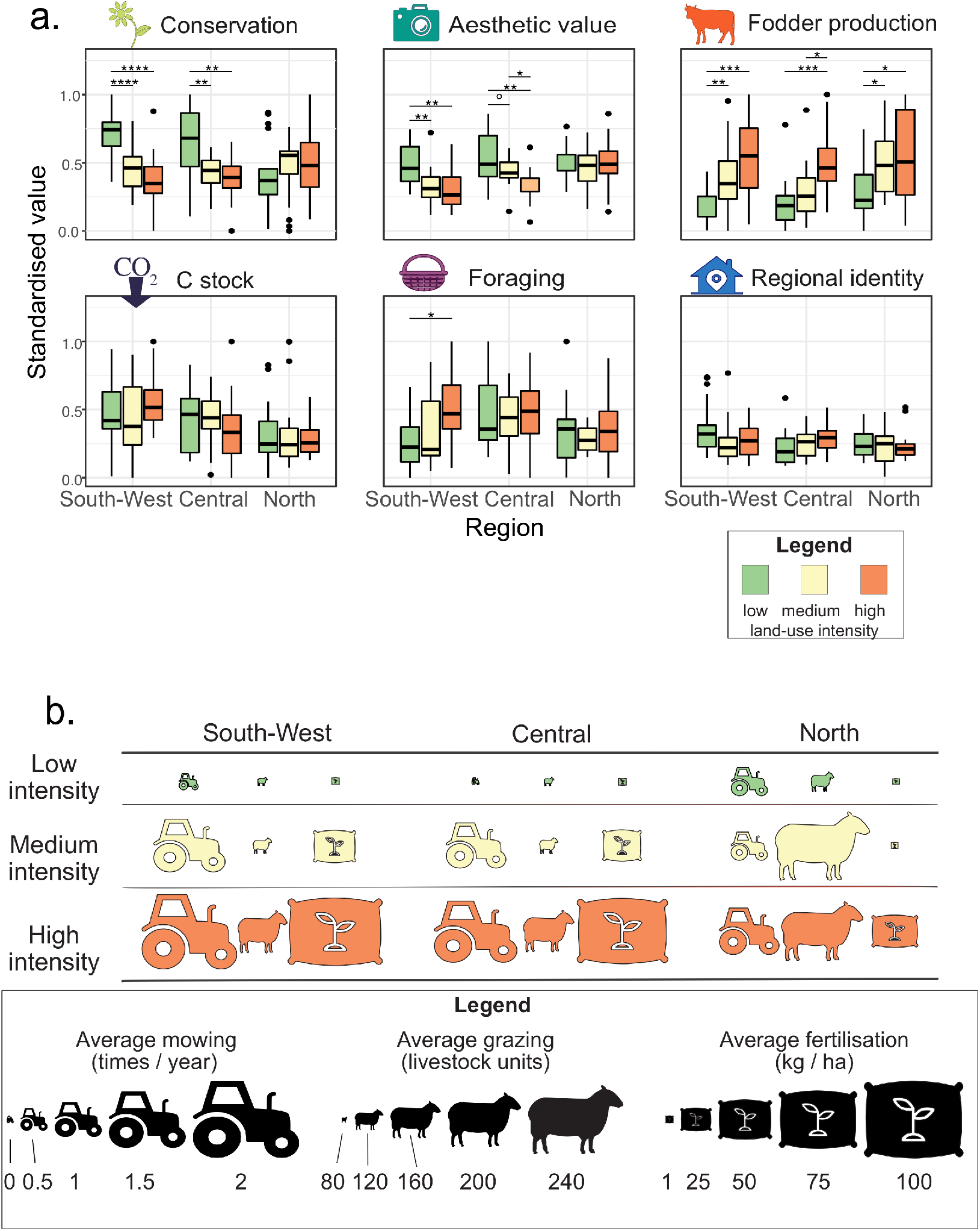
Relationship between ecosystem service supply and land-use intensity across the study regions. a. Variation of ecosystem services supply with land-use intensity. Values shown are calculated at the site level as the average of their component indicators (see Table 1 and supplementary figures). Values were scaled between 0 and 1. Symbols indicate significant differences (ANOVA and pairwise comparisons; **** p < 10^−4^; *** p < 10^−3^; ** p < 10^−2^; * p< 0.05; ° p< 0.1). b. Characterisation of land-use intensity based on mowing, grazing and fertilisation levels in the different regions. The width of the symbols is proportional to average values in each region, in a continuous scale (see Table 2 for full details).

### 2.2 Land-use intensity

Data on site management was collected annually since 2007 from field owners using a questionnaire. We quantified grazing intensity as the number of livestock units × the number of days of grazing (cattle younger than 1 year corresponded to 0.3 livestock units (LU), cattle 1-2 years to 0.6 LU, cattle older than 2 years to 1 LU, sheep and goats younger than 1 year to 0.05 LU, sheep and goats older than 1 year to 0.1 LU, horses younger than 3 years to 0.7 LU, and horses older than 3 years to 1.1 LU; Fischer et al. 2010). Fertilisation intensity was defined as the amount of nitrogen addition, excluding dung inputs from grazing animals (kg N ha^-1^y^-1^), and mowing frequency is the annual number of mowing events. For each site these three land-use intensity (LUI) components were standardised, square-root transformed, summed, and then averaged between 2007 and 2012 to obtain an overall LUI value (Blüthgen et al., 2012). We then classified all sites as low-, medium- or high-intensity based on whether their LUI index (Fig. 1 step 1a) belonged to the lower, middle or top third (0-33%, 33-66%, 66-100% quantiles) of all LUI indices within the considered region (which resulted in a classification equivalent to classifying based on all regions altogether, Fisher test: p < 10^−10^). Confidence intervals for grazing, mowing and fertilisation intensities for each LUI class in the three regions are presented in Table 2. The intensity gradient was mostly driven by fertilisation and cutting frequency in the South-West and Central regions, and by grazing intensity and fertilisation in the North (Fig. 2b), and all three regions span a similar range of LUI values (Table 2).

**Table 1.**
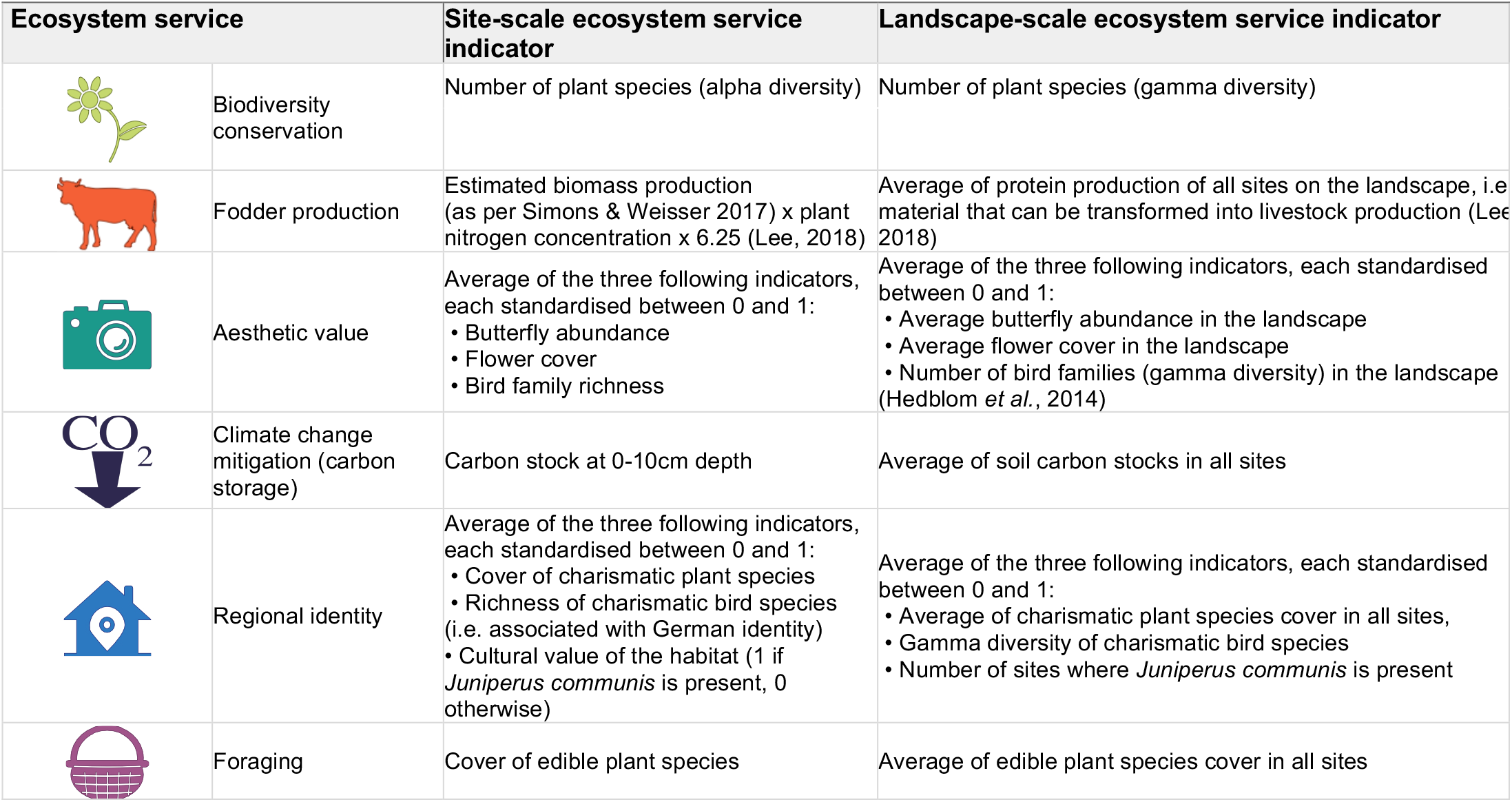
Estimation of ecosystem services from site-scale ecosystem service indicators.

| Ecosystem service |  | Site-scale ecosystem service indicator | Landscape-scale ecosystem service indicator |
| --- | --- | --- | --- |
|  | Biodiversity conservation | Number of plant species (alpha diversity) | Number of plant species (gamma diversity) |
|  | Fodder production | Estimated biomass production (as per Simons & Weisser 2017) x plant nitrogen concentration x 6.25 (Lee, 2018) | Average of protein production of all sites on the landscape, i.e. material that can be transformed into livestock production (Lee 2018) |
|  | Aesthetic value | Average of the three following indicators, each standardised between 0 and 1:<br>• Butterfly abundance<br>• Flower cover<br>• Bird family richness | Average of the three following indicators, each standardised between 0 and 1:<br>• Average butterfly abundance in the landscape<br>• Average flower cover in the landscape<br>• Number of bird families (gamma diversity) in the landscape (Hedblom *et al.*, 2014) |
|  | Climate change mitigation (carbon storage) | Carbon stock at 0-10cm depth | Average of soil carbon stocks in all sites |
|  | Regional identity | Average of the three following indicators, each standardised between 0 and 1:<br>• Cover of charismatic plant species<br>• Richness of charismatic bird species (i.e. associated with German identity)<br>• Cultural value of the habitat (1 if *Juniperus communis* is present, 0 otherwise) | Average of the three following indicators, each standardised between 0 and 1:<br>• Average of charismatic plant species cover in all sites,<br>• Gamma diversity of charismatic bird species<br>• Number of sites where *Juniperus communis* is present |
|  | Foraging | Cover of edible plant species | Average of edible plant species cover in all sites |

**Table 2.** Variation of land-use intensity components. Average (and confidence intervals) for fertilisation, mowing and grazing intensities in each region, for each land-use intensity (LUI) class. 95% confidence intervals were calculated based on the management of individual sites on the period 2007-2012.

|  | LUI class | South-West | Central | North |
| --- | --- | --- | --- | --- |
| LUI index | Low | 1 (0.9-1.2) | 1 (0.9-1.1) | 1.2 (1.1-1.3) |
|  | Medium | 1.7 (1.6-1.7) | 1.7 (1.6-1.8) | 1.5 (1.4-1.5) |
|  | High | 2.2 (2-2.4) | 2.2 (2.1-2.4) | 2.3 (2.1-2.5) |
| Grazing intensity<br>(livestock units.<br>days.ha<sup>-1</sup>) | Low | 82.2 (49.2-115.3) | 86.5 (64.4-108.6) | 103.4 (52.2-154.6) |
|  | Medium | 97.6 (34.5-160.7) | 102.5 (48.3-156.6) | 239.5 (140.9-338.1) |
|  | High | 156.7 (24.6-288.8) | 160.7 (39.6-281.8) | 215.9 (80.4-351.4) |
| Mowing intensity<br>(cut.yr<sup>-1</sup>) | Low | 0.5 (0.1-0.8) | 0.3 (0-0.6) | 0.8 (0.6-1.1) |
|  | Medium | 1.4 (0.9-1.8) | 1.2 (0.9-1.5) | 0.8 (0.4-1.2) |
|  | High | 1.8 (1.3-2.3) | 1.6 (1.2-2) | 1.2 (0.8-1.7) |
| Fertilisation<br>(kg.N.ha<sup>-1</sup>) | Low | 1.1 (-0.6-2.8) | 1.4 (-1.6-4.4) | 0.4 (-0.4-1.2) |
|  | Medium | 38 (23.4-52.7) | 34.9 (18-51.8) | 0.6 (-0.7-2) |
|  | High | 95 (67.4-122.6) | 91.1 (65.4-116.9) | 42.8 (25.8-59.7) |

### 2.3 Ecosystem services priority

Three expert workshops were conducted in autumn 2018 in the three Exploratories, with representatives of numerous pre-selected interest groups. These led to the identification of a full set of 14 interest groups and a list of landscape-level ecosystem services that are demanded regionally. We then restricted the list to services with direct links to final benefits, thus excluding regulating services (e.g. pollination) which underpin the supply of other services (e.g. food production) but do not directly benefit humans. We also excluded water-based services and the production of energy from technology which we weren’t able to cover and both had relatively low weighting. The final list consisted of 12 ecosystem services (Fig. A1). We then conducted a survey across all stakeholder groups in 2019, in which 321 respondents were requested to distribute a maximum of 20 points across all services to quantify their personal priorities. At the beginning of the survey, written consent was requested for the collection and processing of the anonymous personal data; all participants to the survey and workshops could withdraw at any time. More information on the workshops and survey can be found in Peter et al. (in press).

Next, the survey results were subsetted to services provided by grasslands (e.g. removing timber and food crop production), meaning six services were retained: biodiversity conservation, livestock production, aesthetic value, carbon storage, regional identity, and foraging. We then selected, as examples, four stakeholder groups who assigned contrasting priorities to these different services: locals, nature conservation associations, farmers and the tourism sector (126 respondents in total). Priority scores for each service were then normalised by the total number of points attributed to grassland services by each respondent. The priority scores for each group did not vary significantly across regions so we used overall scores.

### 2.4 Ecosystem services supply

We estimated ecosystem service supply from several indicators (Table 1), measured at each of the 150 sites. Biodiversity conservation was based on total plant species richness as plant alpha-diversity at the site level (a good proxy for diversity across multiple trophic levels at these sites, Manning et al., 2015), and gamma-diversity at the landscape level. Fodder production was calculated as total fodder protein production, a common agronomical indicator used in livestock production models, and which better represents livestock production potential than biomass, a large part of which can be indigestible (e.g. Waghorn and Clark, 2004; Herrero, 2013; Lee, 2018). We calculated this indicator from direct measures of grassland aboveground biomass production and shoot protein content. Climate change mitigation was quantified as soil organic carbon stocks in the top 10 cm, as deeper stocks are unlikely to be affected strongly by management actions. For the aesthetic value measure, we integrated direct measures of flower cover, the number of bird families observed and the abundance of butterflies. Regional identity included the species richness (resp. cover) of birds (resp. plants) associated with German identity (see Table B1 and B2), and the foraging service was measured based on the cover of edible plants (see Table B3). Details on the measurements of these different indicators and their aggregation from site to landscape level can be found in the supplementary methods (appendix B).

Before estimating landscape-level services, we imputed missing values for individual indicators using predictive mean matching on the dataset comprising all services (98 out of 1500 values, R mice package v3.13.0; van Buuren & Groothuis-Oudshoorn, 2011). The missing values were mostly for flower cover, and some for butterfly abundance, and were equally distributed among regions and land-use intensities. We also corrected site-level indicator values for environmental covariates (Figure 1, step 1b), thus ensuring differences were caused by management and not covarying factors. Corrected values were the residuals from linear models in which the ecosystem service indicators were the response variable and predictors were: the region, pH, soil depth, sand and clay content, mean annual temperature, mean annual rainfall (see Allan et al. (2015) and Hijmans et al. (2005) for details on these measurements), and a topographic wetness index (see Le Provost et al. 2021 and supplementary methods). To account for a site’s surroundings, we also used the proportion of grassland in a 1km radius as a predictor, as surrounding grassland habitat may act as a source of colonisation for local biodiversity (e.g. Henckel et al., 2015; Le Provost et al., 2017; Tscharntke et al., 2012). Surrounding grassland cover was calculated from 2008 land-cover data that were mapped in QGIS v 3.6 and classified into five broad categories: croplands, grasslands, forests, water bodies, roads and urban areas (Le Provost et al. 2021).

All indicators except gamma diversities were corrected at the site level. This was done to avoid biasing landscape estimates with sites from environmental extremes. These would not be detected when averaging at the landscape level. However, plant and bird species gamma diversity can only be calculated at the landscape level, so were corrected at this scale using the landscape-level averages of the same environmental factors as described above, and also a landscape-level environmental heterogeneity variable. Landscape heterogeneity was calculated as the volume of the convex hull of the selected sites in a PCA that included all environmental variables.

### 2.5 Site-level analyses

We first analysed the relationship between all site-level service indicators and land-use intensity classes. Within each region, we scaled the services between 0 and 1 and fitted ANOVAs with land-use class as the explanatory variable; followed by pairwise mean comparison.

### 2.6 Landscape simulations

An overview of the simulation process is presented in Fig. 1. We conducted the simulations separately for each region, as the relationships between land use and ecosystem services differed strongly between them (Fig. 2).

Landscapes were simulated by randomly drawing sites from the pool of 50 sites from each region (Fig. 1, step 2). Each artificial landscape was composed of ten sites to avoid the high similarity of landscapes composed of more sites. For instance, as there were approximately 16 sites of each intensity class in each region, the most extreme strategies (e.g. 100% low intensity) have only 1 possible landscape composition for landscapes of 16 sites, but 16 possible compositions, which share 90% of their sites for landscapes of 15 sites. For landscapes made up of 10 sites, there are 66 possible landscape compositions that differ in their proportions of low, medium and high intensity sites; ranging from 100% low intensity to 100% medium or high intensity with all possible intermediates (grey dots in the middle triangle of Fig. 1). For each of these compositions, we generated 15 unique artificial landscapes by randomly drawing sites from the regional pool, resulting in 15 × 66 = 990 landscapes. For each simulated landscape, we then calculated landscape-scale ecosystem service indicators, as described above. Simulated landscapes had a combined area of 10 × 50m × 50m sites (i.e. 2.5 ha), but we note that each site is representative of a field ranging from 3 to 148 ha, meaning the results are representative of much larger ‘farm-scale’ landscapes.

Finally, we calculated landscape-scale ecosystem service supply and multifunctionality (Fig. 1, step 3) as described below. We fitted binomial linear models with multifunctionality as the response variable and with a second-degree polynom of the proportions of low and high land-use intensity as explanatory variables (Fig. 1, step 4-5).

### 2.7 Landscape-level ecosystem multifunctionality

The response of all combinations of the six main ecosystem services (i.e. single services, all combinations of 2 to 5 services, and all six services) to landscape composition were investigated. Because trade-offs between services mean that it is unlikely that all services can be maintained at high levels (Bennett et al., 2009; Raudsepp-Hearne et al., 2010; van der Plas et al 2019), managers are often faced with hard choices. To simulate the potential compromises that can be made we therefore generated two contrasting multifunctionality metrics (hereafter multifunctionality scenarios - representing different governance decisions), both of which are multifunctionality measures in which the supply is multiplied by stakeholder priority (Manning et al. 2018). In the first, governors choose to provide a small number of services at high levels, e.g. to meet the needs of a single or few groups to the exclusion of others (hereafter ‘threshold scenario’). In the second, governors opt for a compromise situation in which all services are provided at moderate levels but without any guarantee of them being high (hereafter ‘compromise scenario’). In the threshold scenario, multifunctionality (ranging from 0 to 1) is the proportion of prioritised services that pass a given threshold, here the median value of the service for all landscapes within the considered region. For the compromise scenario we scored multifunctionality as 1 if all services exceeded a 25% threshold (i.e. above the 25% quantile of the service distribution in all landscapes within the region), and 0 otherwise. In addition to these two theoretical scenarios, we also calculated stakeholder-specific multifunctionality values. This was done by scoring each service 1 or 0 depending on whether it passed the 50% threshold (as done in the ‘threshold’ scenario) and averaging the resulting scores, weighted by the relative priority of each service to the stakeholder group (Manning et al. 2018). Priority scores were obtained from points allocation of the social survey (see section 2.3 on Ecosystem services priority).

### 2.8 Dependence of multifunctionality range to the number of services included and to environmental covariates

The response of multifunctionality to landscape composition became increasingly complex as more services were considered (see Results). To understand this complexity, and to identify general rules underlying why optimal landscapes could be identified in some cases but not others, we formed hypotheses and conducted additional analyses. As the objective of these analyses was to find generalities in what affects the responsiveness of multifunctionality to drivers, these were performed upon combined data from all three regions.

Multifunctionality responsiveness is the degree by which multifunctionality responds to landscape composition and it was calculated as the range (2.5% to 97.5% quantiles) of the fitted values of the binomial GLM models described above. Visually, this corresponds to the strength of the colour gradient in the triangle plots presented (Fig. 4 and Fig. 5). While the potential range of multifunctionality is always 1 (i.e. varying from none to all services above the threshold within a landscape), in reality the range of fitted values depends on model fit and the degree of response to landscape composition, or other drivers.

To investigate a hypothesised relationship between multifunctionality responsiveness and the number of ecosystem services included in its calculation, we regressed multifunctionality responsiveness upon the number of ecosystem services included in the landscape-scale assessment (ranging from 1 for individual services to 6 for multifunctionality including all services).

Multifunctionality responsiveness was also hypothesised to depend on the degree of contrast in responses to land-use intensity of the different services included in the assessment. To test this, we estimated slope coefficients of linear regressions between each service and land-use intensity at the site level, and calculated the ‘service response variance’ as the variance of the slope coefficients. We expected that multifunctionality would decrease with the service response variance (van der Plas et al., 2019), but only when services had a strong response to land-use intensity. Thus, we also calculated the ‘service response mean’ as the mean of the absolute values of the slope coefficients. A high service response mean indicates that all services respond strongly to LUI, irrespective of the direction of the response. This service response mean was then classified as ‘high’ (resp. ‘low’) if higher (resp. lower) than the median. We then fitted a linear model in which the range of multifunctionality values was regressed upon the service response variance, the service response mean as a qualitative variable (high or low), and their interaction.

Finally, we tested the hypothesis that multifunctionality was more responsive to land-use intensity when land use played a large role in driving the component functions, relative to other drivers. To do this, we examined the linear relationship between the response of multifunctionality and the relative strength of the LUI effect compared to other environmental covariates. For each single ecosystem service and each region, we quantified the relative strength of the effect of land-use intensity (RS_LUI_) as:

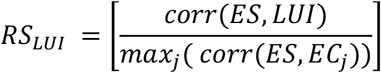

Where *corr* is the correlation, *ES* is the ecosystem service supply value (uncorrected for the environment), *LUI* is the value of land-use intensity, and *EC*_*j*_ are the environmental covariates.

### 2.9 Online tool

Given the sensitivity of the results to the choice of services prioritised and the multifunctionality scenario (see results) we developed an online tool to allow users to investigate which landscape composition best delivers multifunctionality, for any given set of ecosystem service priorities (https://neyret.shinyapps.io/landscape_composition_for_multifunctionality/). This Shiny App (R package shiny v. 1.6, Chang et al. 2021) is based on landscape-scale service supply values pre-calculated for a large range of parameters (see supplementary methods for details on sensitivity analyses) and on a R script similar to the one used for the analyses presented in this manuscript. It allows users to select any combination of parameters (number of sites included, choice of multifunctionality metric, etc.) and service priority scores and to investigate the resulting variations in multifunctionality.

## 3. Results

### 3.1 Relationships between land-use intensity and ecosystem services

At the single-site scale, the optimal land-use intensity for individual services can be easily identified. Across all regions, fodder production consistently increases with land-use intensity while conservation and aesthetic values respond negatively to land-use intensity (Fig. 2). Carbon stocks do not vary with land-use intensity, because environmental variables such as soil texture and mineralogy play a larger role than those of land use within the study regions (Herold et al. 2014). Foraging opportunities and regional identity did not respond consistently to land-use intensity. The trade-offs and synergies between services at the landscape scale (Fig. 3) are consistent with these site-scale results (Fig. 2). Conservation value is synergic with aesthetic value (Pearson’s r = 0.28 for all regions, P < 0.001) but both display a trade-off with fodder production (respectively r = - 0.21 and r = - 0.41, P < 0.001). The other services do not show any consistent correlation.

**Figure 3.**
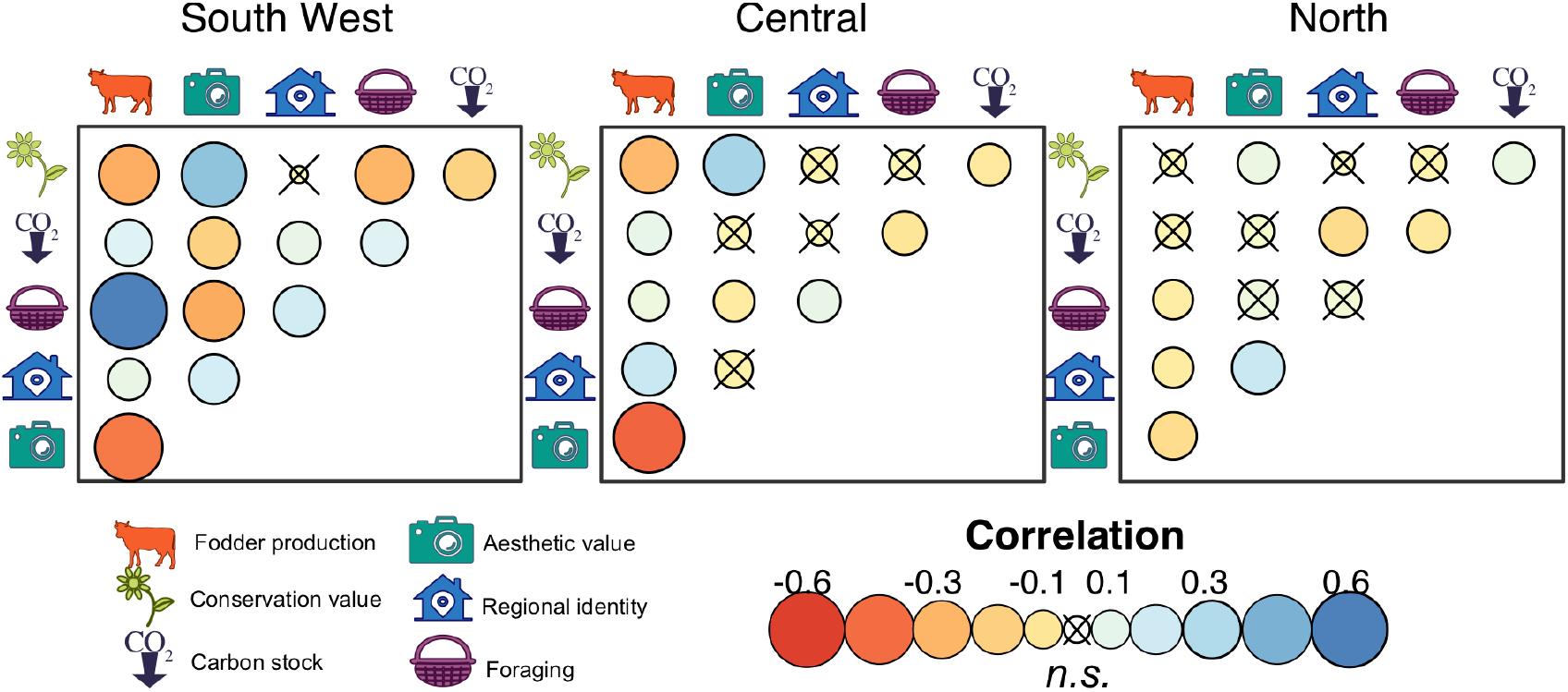
Trade-offs between landscape-scale ecosystem services. The colour and size of the circles denote the strength of the correlation between pairs of variables, within each region. Crosses indicate no significant correlations at 5% (Holm correction for multiple testing).

### 3.2 Optimal land-use allocation at the landscape scale

At the landscape scale, effective landscape strategies can be identified where only a few services are desired, but optimisation becomes increasingly difficult as more services are considered. The optimal land-use allocation pattern also depends strongly on whether achieving moderate levels of all services, or high levels of a few, is the priority. This sensitivity is best explored in the online tool (https://neyret.shinyapps.io/landscape_composition_for_multifunctionality/), which allows users to investigate the best management strategy given any set of ecosystem service priorities and the impact of land-use changes from current conditions. In the text below we highlight a few of the possible combinations of this parameter space, and demonstrate the sensitivity of multifunctionality to multiple factors. We then illustrate our results using data collected from four of the main stakeholder groups of the three study regions: farmers, conservationists, locals and the tourism sector.

In the first set of examples, we present the ‘threshold’ scenario in which land governors consider a landscape that provides high levels of some services, potentially to the exclusion of some others. Here we find that for individual services, the optimal landscape composition is predictable and consistent with the site-level results, i.e. that when services are significantly affected by land-use intensity, the highest service values are found in homogeneous landscapes composed of sites with land-use intensities favouring that particular service (Fig. 4a-c). The optimal landscape composition when two ecosystem services are considered depends on whether these services have consistent or contrasting responses to land-use intensity. When the two services are synergic, they behave as a single service and optimal landscape composition is found at the common optimum of the two services. For example, a clear optimum can be found for conservation and aesthetic value (Fig. 4i). In contrast, if the two services respond contrastingly to land-use intensity, then whether an optimum could be found depends on the form and strength of their relationship with land-use intensity. For example, a common objective of landscape management is to combine food production with biodiversity conservation (Phalan et al. 2011). As there is a strong trade-off between these services (Fig. 3), only a partial optimum with high levels of both services can be found (Fig. 4g), with the landscape composition delivering this depending on regional differences in the response of services to land-use intensity (see Fig. A2 for details), and the relative responsiveness of the services considered to land-use intensity. For three or more services (Fig. 4j-l) the identification of a clear optimal land-use strategy becomes even more challenging. In these cases, multifunctionality varies very little across the full range of landscape composition (maximum R^2^ 15%, and often < 10%, Fig. A2), with relatively uniform multifunctionality values of about 50%, regardless of the landscape composition.

**Figure 4.**
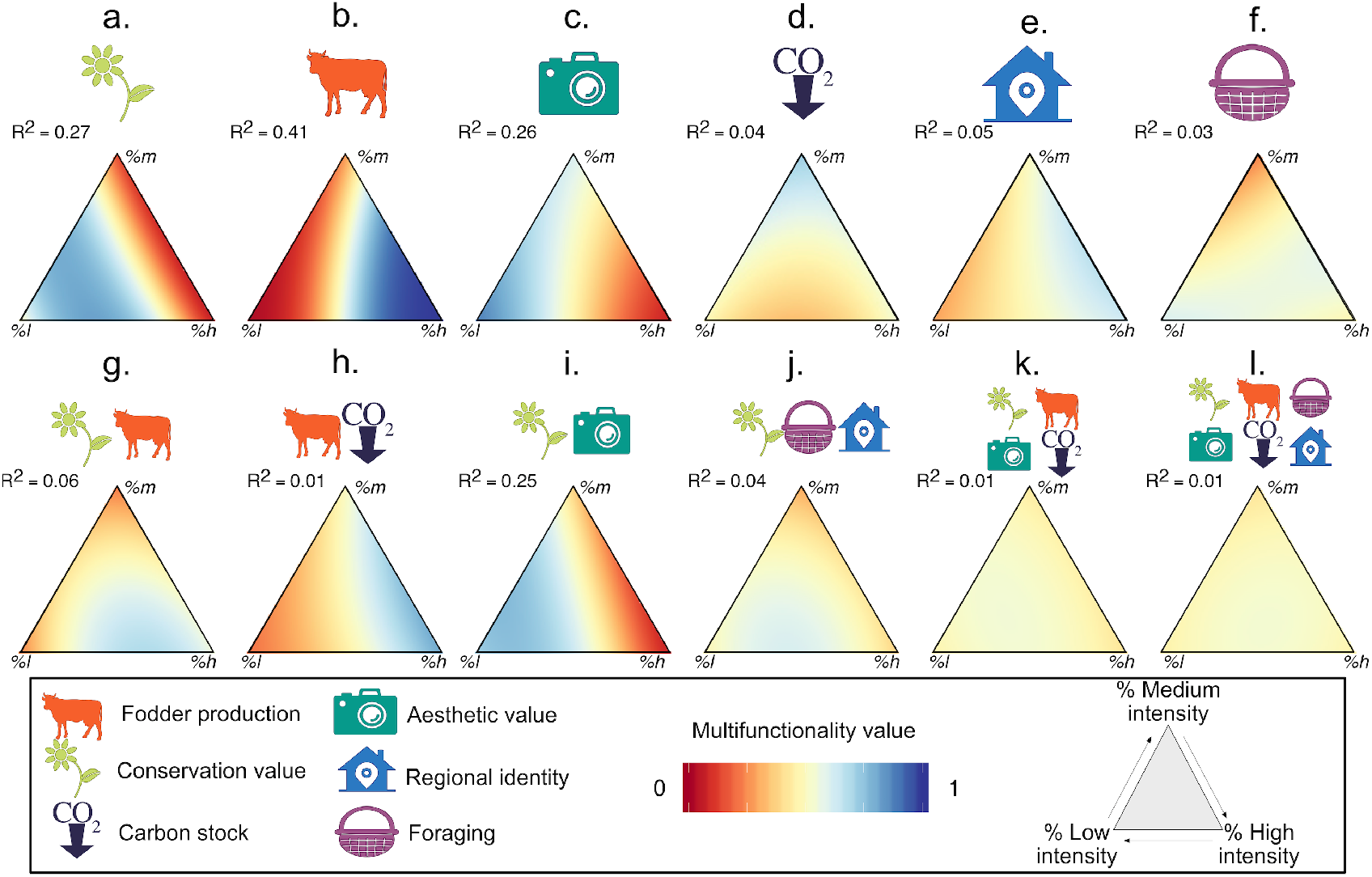
Dependency of overall ecosystem service supply on the prioritized services and landscape composition. Landscape composition is presented as proportions of low, medium and high-intensity sites, for selected combinations of ecosystem services in the Central region of the Exploratories. For single ecosystem services (top row), the service supply corresponds to the probability of the given service being above the median. For combinations of multiple services (bottom row), multifunctionality is the proportion of services above the median, averaged across multiple simulated landscapes. A broader set of service combinations in all regions can be found in Supplementary Figure A2. R^2^ values were calculated from generalised linear models (see Methods).

In the ‘compromise’ scenario, land governors consider a landscape multifunctional if it balances the demands of different stakeholder groups, ensuring moderate, but not necessarily high, level of all services. This gives very different results in comparison to the ‘threshold’ scenario (for selected service combinations see Fig. 5). The two scenarios give similar results when services are synergic (e.g. Fig. 5b), but when the considered services display a trade-off it is easier to identify successful land-use strategies in the ‘compromise’ scenario, especially when there are only two services (Fig. 5a). In this case ‘compromise’ multifunctionality is highest in landscapes composed of both high- and low-intensity sites, and with few medium-intensity sites, i.e. broadly similar to a land-sparing strategy, and consistent with our original hypothesis. When multiple services are considered (Fig. 5d-f), the variation in multifunctionality values across the different land-use strategies also tends to be higher in the ‘compromise’ than the ‘threshold’ scenario. For example, see the flat response of multifunctionality to land use in the threshold scenario (values all ∼0.5, Figure 5f) compared to the wider range of the compromise scenario (values 0-0.4). This indicates that in the compromise scenario, an optimum can be identified even though maximum multifunctionality is low.

**Figure 5.**
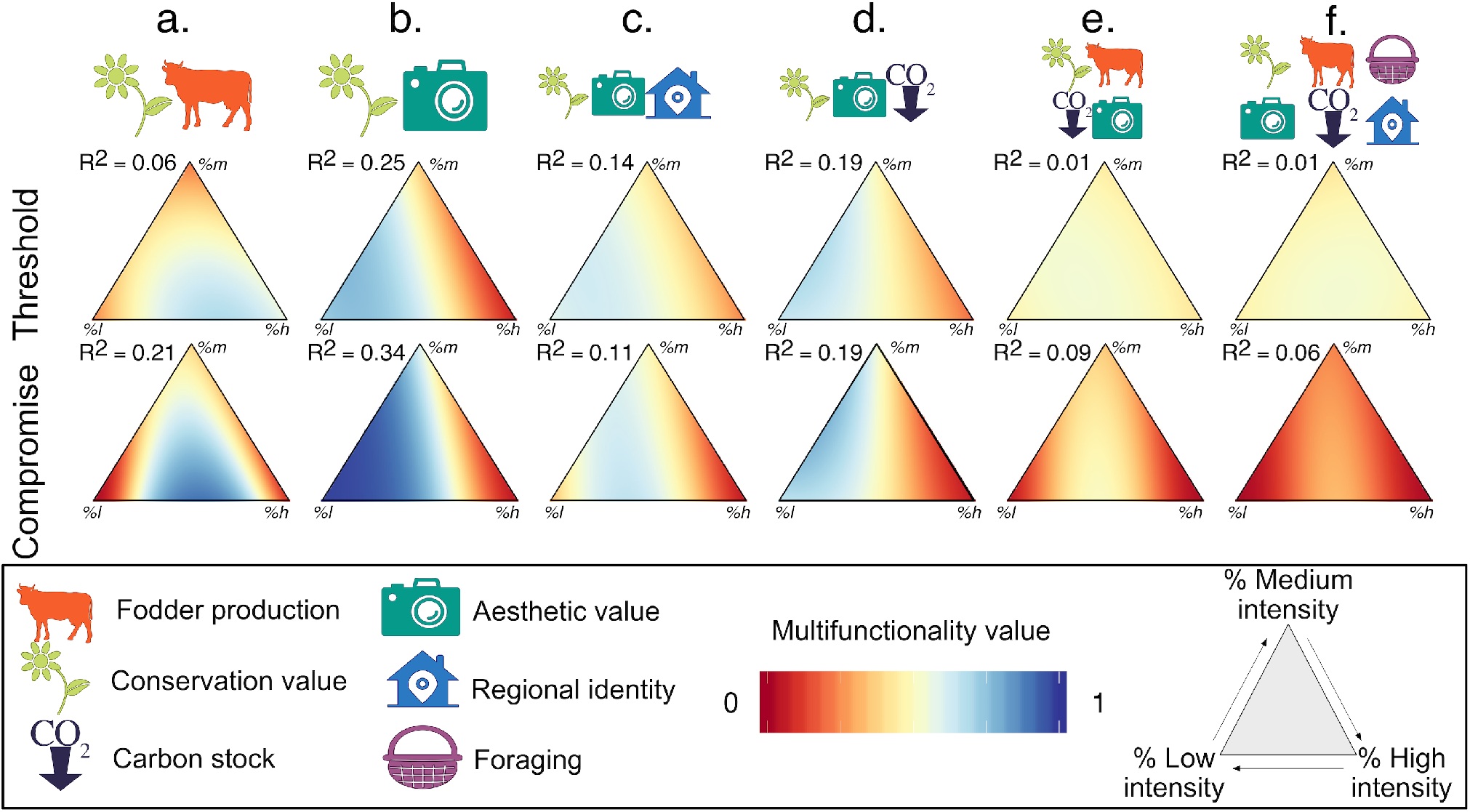
Dependency of multifunctionality on stakeholder demand patterns, as represented by ‘threshold’ and ‘compromise’ scenarios. Values also depend on landscape composition (in proportions of low, medium and high-intensity sites). In the threshold scenario, multifunctionality is calculated as the proportion of services above the median (top row, partly repeated from Fig. 4). In the compromise scenario, multifunctionality equals 1 if all services are above the 25^th^ quantile, and 0 otherwise (bottom row). In both cases, the values are averaged across multiple simulated landscapes, hence the continuous values. R^2^ values were calculated from generalised linear models (see Methods). Only data from the Central region and certain service combinations are presented, other service combinations and regions can be found in Supplementary Figures A2 and C7.

### 3.3 Optimal land-use for contrasting stakeholder groups

Different stakeholder groups had contrasting service demands, but all groups allocated points to all services (Fig. A1). This demand for multiple services resulted in relatively weak responses of overall multifunctionality to landscape composition (Fig. A3). Consistent with the results described above, the optimal landscape also depended on the relative priority of different services. For instance, farmers reported a clear priority for productivity (40% of the total priority score for grassland services) and so find their optimum in landscapes composed of mostly high-intensity sites. In contrast, nature conservation associations prioritised biodiversity (39%) and so found their optimum in landscapes dominated by low-intensity sites. Tourists and locals weighted services more equally (e.g. 26% conservation, 22% aesthetic value, 20% regional identity for tourists) meaning their optimum landscape compositions were less easy to identify, although an optimum was usually found for intermediate to high proportions of low-intensity sites.

### 3.4 Identifying the drivers of multifunctionality

To explore why optimal landscape strategies cannot always be identified when multiple services are prioritised, we tested several hypotheses. Hypothesis 1, some services are primarily driven by environmental drivers (e.g. climate and underlying geology) and thus respond weakly to changes in landscape composition. Hypothesis 2, if services respond contrastingly to land-use intensity (i.e. trade-off), then their respective contributions to multifunctionality counteract each other, resulting in a flat response of multifunctionality to changes in landscape composition. Hypothesis 3, increasing the number of services demanded increases the chance that service responses will contrast, and also aggregates increasing amounts of variation, thus weakening the response of multifunctionality to landscape composition.

Hypothesis 1 was supported: in the threshold scenario, the responsiveness of multifunctionality to landscape composition (defined as the range of fitted multifunctionality values, visualised as “colourfulness” in Fig. 4-5; see Methods) increased when land-use intensity had a relatively large effect on the services compared to other environmental drivers (Fig. 6a, P < 10^−3^, R^2^ = 50%). Hypotheses 2 and 3 were also supported; variation in multifunctionality decreased with increasing variance in the response of services to land-use intensity, at least when the service response was strong (service response mean above the median) (Fig. 6c, P < 10^−3^, R^2^ = 11%), and with increases in the numbers of services included in the analysis (Fig. 6b, P < 10^−3^. R^2^ = 21%). Most of these relationships were also found in the compromise scenario (Table C1).

**Figure 6.**
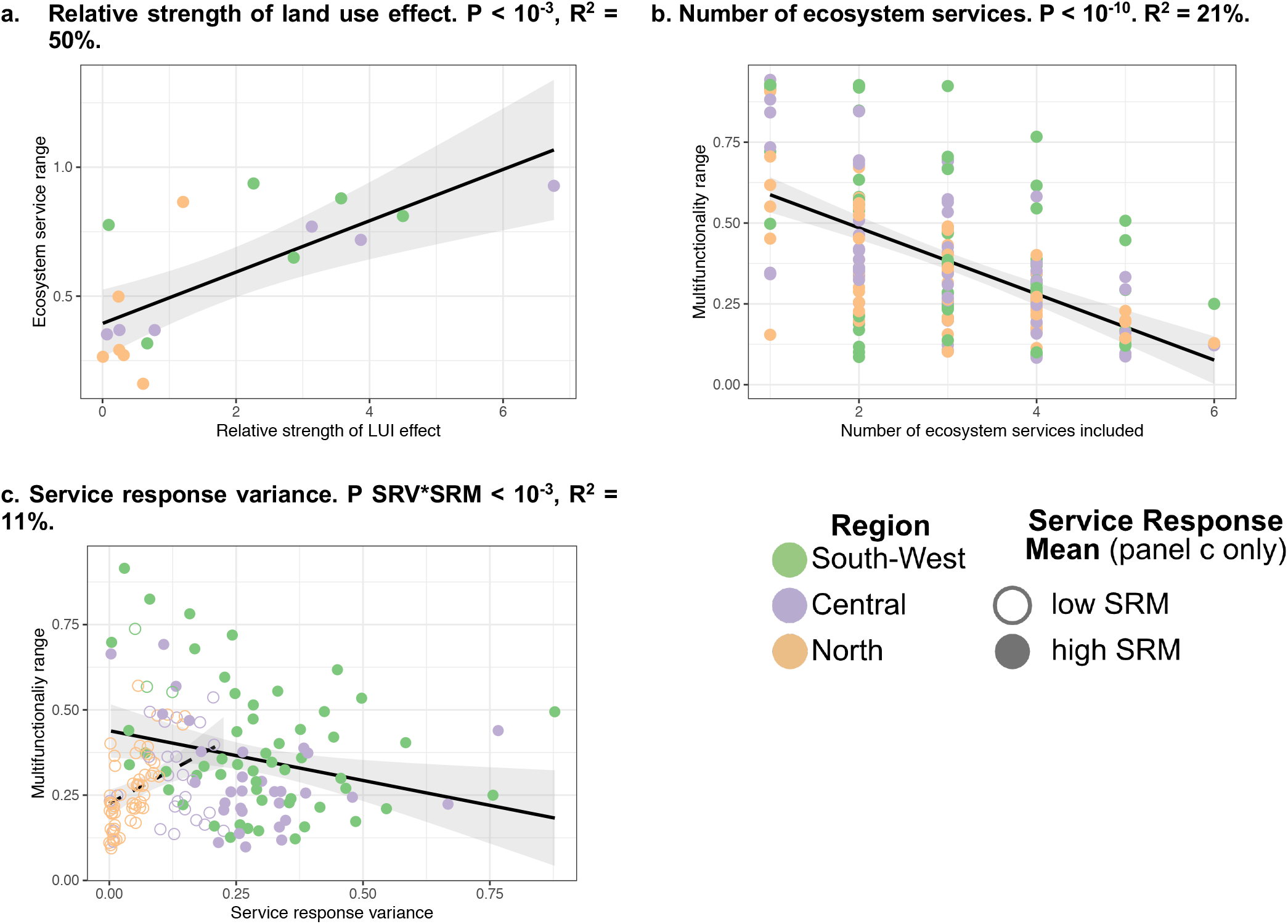
Factors affecting the responsiveness of overall ecosystem service supply to landscape composition. Figures show the responsiveness of single service supply or multifunctionality (range between 5% and 95% quantiles of the predicted values) to landscape composition depending on (a) the strength of individual ecosystem services’ response to landscape composition relative to the effects of land use and environmental covariates (b) the number of ecosystem services included in the calculation of multifunctionality (all possible combinations, of 1 to 6 services) and (c) the service-response variance among the included ecosystem services (all possible combinations); P SRV*SRM indicates p-value for the interaction between service response variance and the service response mean (low (resp. high) is lower (resp. higher) than the median). Each dot represents individual services (a), or one combination of services (b, c), per region. The lines and shaded areas show the prediction of a linear model (and confidence intervals), with multifunctionality range as the response and the considered factor as the explanatory variable. In panel c, filled dots and solid lines show high mean response; and empty dots with dashed lines low mean response.

### 3.5 Additional analyses

In addition to the main cases presented here, we identified several other sensitivities including additional metrics for multifunctionality calculation at the landscape level, the number of sites included in each landscape, and the use of raw data instead of that corrected for environmental variation. We encourage readers to explore these sensitivities in the app, although the corresponding figures and specificities are also presented extensively in the supplementary information (Supplementary Fig. C3 to C15). In addition, we also investigated the response of multifunctionality to the mean and coefficient of variation of land-use intensity at the landscape level. The results were similar to those described here, in that unless the services were synergic, no clear optimum could be found when several services were demanded (Fig. C17).

## 4. Discussion

While the land-sharing or -sparing debate has aided our understanding of the trade-offs between commodity production and conservation (Phalan, 2018) we show that strategies that are broadly comparable to these cannot provide high multifunctionality in grassland landscapes, if high levels of multiple ecosystem services are desired. In particular, while our approach allows stakeholders to identify an optimal landscape composition for any given set of priorities, demand for multiple ecosystem services typically led to low to moderate multifunctionality that hardly varied with landscape composition. This indicates that there is no landscape composition in which supply can meet the demand for all services. We predict that this difficulty in achieving high multifunctionality is general to many ecosystems and landscapes, as the presence of other drivers and weak and negative correlations between services are commonplace (Bennett et al., 2009). Various studies have advocated for the consideration of more complex strategies for balancing commodity production with conservation (Bennett, 2017; Butsic & Kuemmerle, 2015; Chan et al. 2006; Fischer et al., 2014; Phalan et al., 2011, Simons and Weisser 2017). By employing a rigorous approach based on direct, in-field measurements of ecosystem service indicators, we further show that considering not only trade-offs and synergies between ecosystem services, but also information describing the ecosystem service priorities of stakeholders, helps identify land management options that have greater precision and relevance to land users. The approach presented also allows the potential causes of land-use conflicts to be identified, as it can assess whether low multifunctionality is caused by trade-offs in the supply of ecosystem services, or unrealistic and incompatible demands on the ecosystem by stakeholders. Furthermore, the impact of any given land-use change, e.g. deintensification of 50% of medium intensity grasslands, on stakeholders with different priorities can be estimated and accounted for in decision making.

In our study system, ecosystem services showed contrasting responses to land-use intensity, including the commonly observed trade-off between production and biodiversity or cultural services (Allan et al., 2015; Cordingley et al., 2016; Lavorel et al., 2011; Raudsepp-Hearne et al., 2010). Understanding contrasting responses of ecosystem services to land management is fundamental to identifying landscape-level strategies. Here, we show that strong management-driven trade-offs preclude multifunctionality when high levels of services are required. As a result, even complex landscape strategies may fail to deliver high levels of multiple ecosystem services (Allan et al., 2015) and landscape management is likely to require “hard choices” (Cordingley et al., 2016; Slade et al., 2017) regarding which services to prioritise, and which are secondary. A social survey of stakeholders in our study system showed that all stakeholders demanded multifunctionality (supporting previous results, e.g. Hölting et al. 2020). Although it is possible to find a landscape optimum when stakeholders clearly prioritise one service (e.g., farmers and livestock production), the multifunctionality demanded makes landscape optimisation difficult even for a single stakeholder group, as multiple services that do not synergise were often given equally high priority.

Although high levels of all services may be unattainable, we show that it is possible to provide limited levels of multiple services by combining sites at low and high intensities, a strategy broadly similar to land-sparing. In this respect, our results show that the optimal strategy depends heavily on the levels of service provision landscape managers are willing to accept. While different stakeholders favour different sets of services, landscape-level governors are faced with a difficult choice: create a landscape with a few services at high value, which will create winners and losers among stakeholder groups, or one that minimises the trade-offs among services so that all are present at moderate levels, meaning that all stakeholder groups must accept sub-optimal levels of ecosystem services.

While advancing on previous studies by incorporating multiple services, we acknowledge that our approach to identifying optimal landscape strategies is simple and ignores much of the complexity found in natural systems. First, land-use decisions are also affected by the relationship between ecosystem service supply and the amount of benefit provided: while we considered the supply-benefit relationship for all services as behaving in a threshold manner, this is clearly a simplification. More accurate and service-specific representation of the relationships between supply and benefit (e.g. Linders et al. 2021) could provide additional insights into which landscape compositions best deliver multifunctionality. For instance, at very low intensity fodder production might not be profitable enough for farmers, resulting in no benefit despite some supply, and with a risk of pasture abandonment. While tailored supply-benefit relationships have been rarely used in multifunctionality metrics due to the difficulty in defining their shape, their integration could greatly improve model predictions (Manning et al. 2018, Linders et al. 2021). Both the prioritisation of services by stakeholders, and supply-benefit relationships, are also likely to change in time and space (Boesing et al. 2020), e.g. as market forces and society change. Incorporating this temporal dimension remains a challenge for future multifunctionality research.

An aspect highlighted by our study is the difficulty in identifying land management strategies when ecosystem services respond to multiple drivers. Additional drivers can be either anthropogenic (e.g. other land-use changes, overexploitation, Carpenter et al., (2009)) or environmental (e.g. soil (Adhikari & Hartemink, 2016), climate, or elevation (Lavorel et al., 2011)). Failing to account for these drivers can obscure the relationship between land-use composition and multifunctionality. In this study, the use of real-plot data at least partly integrates this inherent variability, which also explains why some trends are not as clear-cut as those that could be obtained from models using fixed values. Environmental drivers will differ in their effect on different services, and so can modify their trade-offs (Le Clec’h et al., 2019). Therefore, the development of strategies to achieve landscape multifunctionality also needs to be informed by regional knowledge (Butsic & Kuemmerle, 2015; Le Clec’h et al., 2019). For instance, in our analysis the Northern region responded very differently to the other two regions. This was due to regional specificities, such as its uniformly low plant diversity and the association of low-intensity sites with organic soils, which shifted the optimal landscape compositions to different regions of the triangular space compared to the other regions (Fig. A2). Despite this context-dependency, we expect that our study areas are representative of their regions both in terms of environmental conditions and management, making our results broadly applicable to Germany.

In addition to local drivers, the delivery of many ecosystem services depends on the movement of matter or organisms among landscape units (Mitchell et al., 2014). For instance, pollination, water quality, or pest and disease control are affected by landscape complexity, fragmentation and surrounding land uses (Duarte et al., 2018). Accordingly, we advocate the incorporation of spatial interactions between landscape units (Lindborg et al., 2017) into future models, elements which may modify and expand upon the conclusions presented here.

Our system consists of only one land-use type and does not include unmanaged land. This makes it only broadly comparable to the land-sparing and -sharing strategies, which typically integrate more diverse land-use types and also aspects relating to their spatial organisation in a landscape (e.g., land sharing strategies may lead to a patchwork of intensively farmed and semi-natural landscape units, in which overall service provision may be influenced by their configuration). However, we argue that the approach presented here could be extended to many different land-use and management regimes, provided that appropriate data on services and drivers is available. Steps must also be taken to ensure that insights from such studies are in a format that can be communicated effectively to land managers. For instance, we argue that presenting proportions of land in a number of land-use categories is more easily transferable than indices of land-use intensity heterogeneity. Strategies for knowledge transfer also need to be developed. We suggest that online tools like the one presented here provide a useful demonstration tool for communicating land-use options to land managers and policymakers, as they could be used to explore options, understand the causes of conflicts and trigger discussions, thus helping to support decision-making among different groups of stakeholders. However, the full application of findings such as those presented here also requires the existence of structures that aim to identify landscape strategies and operationalise them at a community level, such as the ‘landscape approach’ (DeFries & Rosenzweig, 2010; Sayer et al., 2013). This aims to balance competing land-use priorities to promote environmental conservation and human well-being based on a participatory approach (e.g. the African Forest Landscape Restoration Initiative). Government and corporate policies can also implement such strategies, e.g. via agri-environment schemes that may guide the allocation of different land-use types or land-use intensities to different parts of the landscape (Whittingham, 2011). We suggest that demonstrating the impacts of different management options via apps such as that presented here, can foster understanding and aid decision making in both of these settings.

Overall, this study shows that landscape strategies are highly sensitive to the identity of the services desired and the type of multifunctionality demanded by stakeholders, making participatory approaches to the development of land management strategies essential. When high levels of all services are required, we show that optimising landscape composition is usually possible for two services. However, when there are strong trade-offs among services or significant effects of other environmental drivers, winning management options become increasingly hard to identify unless stakeholders are willing to accept moderate service levels, which can be delivered by strategies akin to land sparing. Across the world, landscapes are increasingly required to provide a wide range of services. This study stresses the need for both theoretical studies and applied social and ecological research into which services are required, at what scale, and how they are affected by environmental drivers. Such knowledge is essential if we are to identify land-use strategies that minimise conflict between stakeholders, and promote the sustainable use of all ecosystem services.

## Supporting information

Sensitivity analyses

## 5. Acknowledgements

We thank Steffen Boch for his help in designing the list of edible and culturally important plants. We thank the managers of the three Exploratories Konstans Wells, Swen Renner, Kirsten Reichel-Jung, Sonja Gockel, Kerstin Wiesner, Katrin Lorenzen, Andreas Hemp, Martin Gorke and all former managers for their work in maintaining the plot and project infrastructure; Simone Pfeiffer, Maren Gleisberg and Christiane Fischer for giving support through the central office, Jens Nieschulze and Michael Owonibi for managing the central data base, and Markus Fischer, Eduard Linsenmair, Dominik Hessenmöller, Daniel Prati, Ingo Schöning, François Buscot, Ernst-Detlef Schulze, Wolfgang W. Weisser and the late Elisabeth Kalko for their role in setting up the Biodiversity Exploratories project.

The work has been funded by the DFG Priority Program 1374 “Infrastructure-Biodiversity-Exploratories”. Fieldwork permits were issued by the responsible state environmental offices of Baden-Württemberg, Thüringen, and Brandenburg.

## 6. Data accessibility

The code to reproduce the results presented in this paper and the Shiny app are available on GitHub/Zenodo 10.5281/zenodo.5521456. Pre calculated, landscape-level data is available is available from the Biodiversity Exploratories information sytem (Bexis) (https://doi.org/10.25829/bexis.31094-15).

## Notes

https://doi.org/10.5281/zenodo.5521457

https://doi.org/10.25829/bexis.31094-15

### Competing Interest Statement

The authors have declared no competing interest.

### Summary of Updates

Add dataset DOI Correct section numbering

