## Additional information for "Assessing the impact of grassland management on landscape multifunctionality"

### Appendix A: Additional information

Neyret M., Fischer M., Allan E., Hölzel N., Klaus V. H., Kleinebecker T., Krauss J., Le Provost G., Peter. S., Schenk N., Simons N.K., van der Plas F., Binkenstein J., Börschig C., Jung K., Prati D., Schäfer D., Schäfer M., Schöning I., Schrumpf M., Tschapka M., Westphal C. & Manning P.

**Table A1 Variation in the ecosystem service measures and their component service indicators**, with the land-use intensity class in each region. Values were first corrected for the environment (see Methods). Different letters indicate differences significant at 5%.

| **Ecosystem services** or *service indicator* | Land-use intensity | South-West | Central | North |
|---|---|---|---|---|
| **Livestock production** | low | -66.45 +/- 18.44 [a] | -37.2 +/- 14.36 [a] | -34.97 +/- 14.71 [a] |
| | medium | 9.9 +/- 19 [b] | -10.13 +/- 14.8 [a] | 11.09 +/- 15.17 [ab] |
| | high | 57.12 +/- 18.44 [b] | 46.73 +/- 14.36 [b] | 24.53 +/- 14.71 [b] |
| *Biomass production* | low | -33.44 +/- 7.04 [a] | -20.06 +/- 5.84 [a] | -15.8 +/- 5.16 [a] |
| | medium | 5.69 +/- 7.25 [b] | -4.54 +/- 6.02 [a] | 5.56 +/- 5.31 [b] |
| | high | 28.09 +/- 7.04 [b] | 24.33 +/- 5.84 [b] | 10.57 +/- 5.16 [b] |
| *N content* | low | -0.01 +/- 0.08 [a] | -0.1 +/- 0.07 [a] | 0.07 +/- 0.13 [a] |
| | medium | 0.01 +/- 0.08 [a] | 0.02 +/- 0.08 [a] | -0.03 +/- 0.13 [a] |
| | high | -0.01 +/- 0.08 [a] | -0.02 +/- 0.07 [a] | 0.07 +/- 0.13 [a] |
| **Aesthetic** | low | 0.51 +/- 0.03 [b] | 0.49 +/- 0.04 [b] | 0.43 +/- 0.02 [a] |
| | medium | 0.37 +/- 0.03 [a] | 0.41 +/- 0.04 [ab] | 0.4 +/- 0.03 [a] |
| | high | 0.33 +/- 0.03 [a] | 0.33 +/- 0.04 [a] | 0.41 +/- 0.02 [a] |
| *Flower cover (sqrt)* | low | 0.17 +/- 0.2 [a] | 0.29 +/- 0.24 [a] | -0.06 +/- 0.15 [a] |
| | medium | -0.25 +/- 0.21 [a] | -0.02 +/- 0.25 [a] | 0 +/- 0.16 [a] |
| | high | 0.06 +/- 0.2 [a] | -0.27 +/- 0.24 [a] | 0.07 +/- 0.15 [a] |
| *Butterfly abundance (sqrt)* | low | 2.75 +/- 0.73 [b] | 1.32 +/- 0.77 [b] | 0.56 +/- 0.46 [a] |
| | medium | -0.48 +/- 0.76 [a] | 0.28 +/- 0.79 [ab] | -0.58 +/- 0.47 [a] |
| | high | -2.7 +/- 0.73 [a] | -1.36 +/- 0.77 [a] | 0.16 +/- 0.46 [a] |
| *Bird family richness* | low | 0.95 +/- 0.54 [a] | 0.84 +/- 0.72 [a] | 0.49 +/- 0.5 [a] |
| | medium | -0.24 +/- 0.56 [a] | 0.28 +/- 0.74 [a] | -0.03 +/- 0.52 [a] |
| | high | -0.88 +/- 0.54 [a] | -0.99 +/- 0.72 [a] | -0.42 +/- 0.5 [a] |
| **Conservation** *(plant richness)* | low | 9.24 +/- 1.97 [b] | 10.13 +/- 2.64 [b] | -0.71 +/- 1 [a] |
| | medium | -3.04 +/- 2.03 [a] | -3.77 +/- 2.72 [a] | -0.05 +/- 1.03 [a] |
| | high | -6.38 +/- 1.97 [a] | -6.58 +/- 2.64 [a] | 0.76 +/- 1 [a] |
| **C stock** | low | -0.04 +/- 0.22 [a] | -0.02 +/- 0.24 [a] | -0.02 +/- 0.56 [a] |
| | medium | -0.27 +/- 0.22 [a] | 0.11 +/- 0.25 [a] | 0.27 +/- 0.58 [a] |
| | high | 0.3 +/- 0.22 [a] | -0.08 +/- 0.24 [a] | -0.23 +/- 0.56 [a] |
| **Foraging** *(cover edible plants)* | low | -25.63 +/- 12.12 [a] | 4.06 +/- 12.12 [a] | 11.4 +/- 22.52 [a] |
| | medium | -2.57 +/- 12.5 [ab] | -3.59 +/- 12.49 [a] | -18.51 +/- 23.22 [a] |
| | high | 23.98 +/- 12.12 [b] | 2.21 +/- 12.12 [a] | 7.2 +/- 22.52 [a] |
| **Regional ID** | low | 0.36 +/- 0.04 [a] | 0.24 +/- 0.03 [a] | 0.26 +/- 0.03 [a] |
| | medium | 0.25 +/- 0.04 [a] | 0.27 +/- 0.03 [a] | 0.25 +/- 0.03 [a] |
| | high | 0.27 +/- 0.04 [a] | 0.31 +/- 0.03 [a] | 0.24 +/- 0.03 [a] |
| *Juniperus grasslands* | low | 0.24 +/- 0.07 [a] | 0 | 0 |
| | medium | 0.06 +/- 0.07 [a] | 0 | 0 |
| | high | 0 +/- 0.07 [a] | 0 | 0 |
| *Cover cultural plants* | low | -13.61 +/- 14.88 [a] | -23.4 +/- 15.2 [a] | -11.39 +/- 18.84 [a] |
| | medium | -17.34 +/- 15.34 [a] | -0.68 +/- 15.67 [ab] | -1.86 +/- 19.42 [a] |
| | high | 27.06 +/- 14.88 [a] | 34.08 +/- 15.2 [b] | 5.98 +/- 18.84 [a] |
| *Richness cultural birds* | low | 0.51 +/- 0.31 [a] | 0.18 +/- 0.39 [a] | 0.17 +/- 0.27 [a] |
| | medium | -0.15 +/- 0.32 [a] | 0.16 +/- 0.4 [a] | -0.23 +/- 0.28 [a] |
| | high | -0.47 +/- 0.31 [a] | 0.1 +/- 0.39 [a] | -0.28 +/- 0.27 [a] |

**Figure A 1. Stakeholder groups' ecosystem preferences.** a. Mean demand score for all considered services, per stakeholder group (coloured dots) or all groups considered (crosses). We then retained for the analysis only the six main services that can be delivered by grasslands (red box). Four stakeholder groups are presented as an example in this paper; they are shown with full dots while the other groups are shown with empty circles. b. Mean priority scores for the services included in the analysis of the current paper, for the selected stakeholder group; shown as the proportion of points attributed to each service relative to the total number of points attributed to the six services.

a.

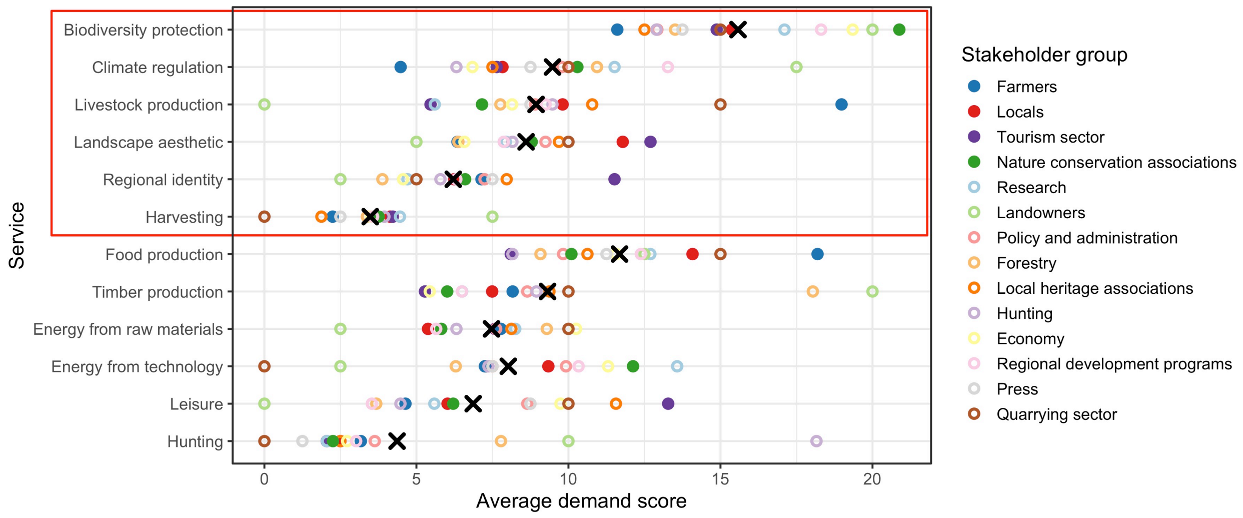


b.

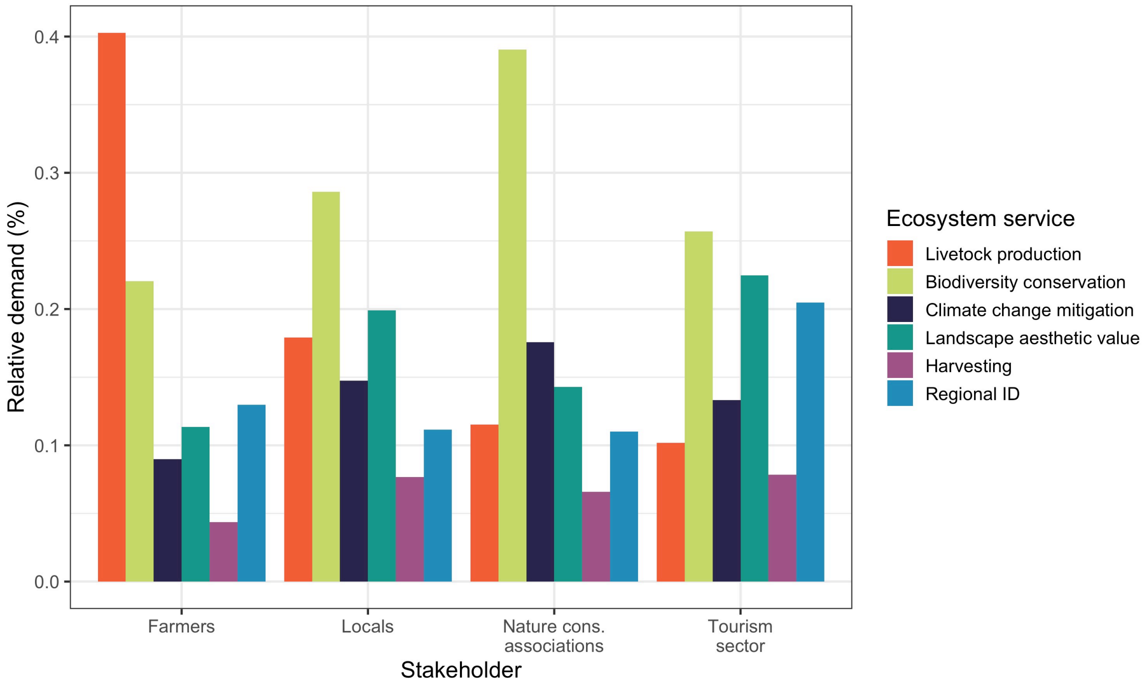

**Figure A2** Estimated multifunctionality values depending on landscape composition (in proportions of low, medium and high-intensity sites). **This figure shows a wider selection of the service combinations for the 'threshold' approach shown in Fig. 4.**

For single ecosystem services (top row), the value presented corresponds to the probability of the given service to be above the median. For combinations of multiple services (middle and bottom rows), multifunctionality is the expected proportion of services above the median. Blue indicates higher multifunctionality values, orange lower.

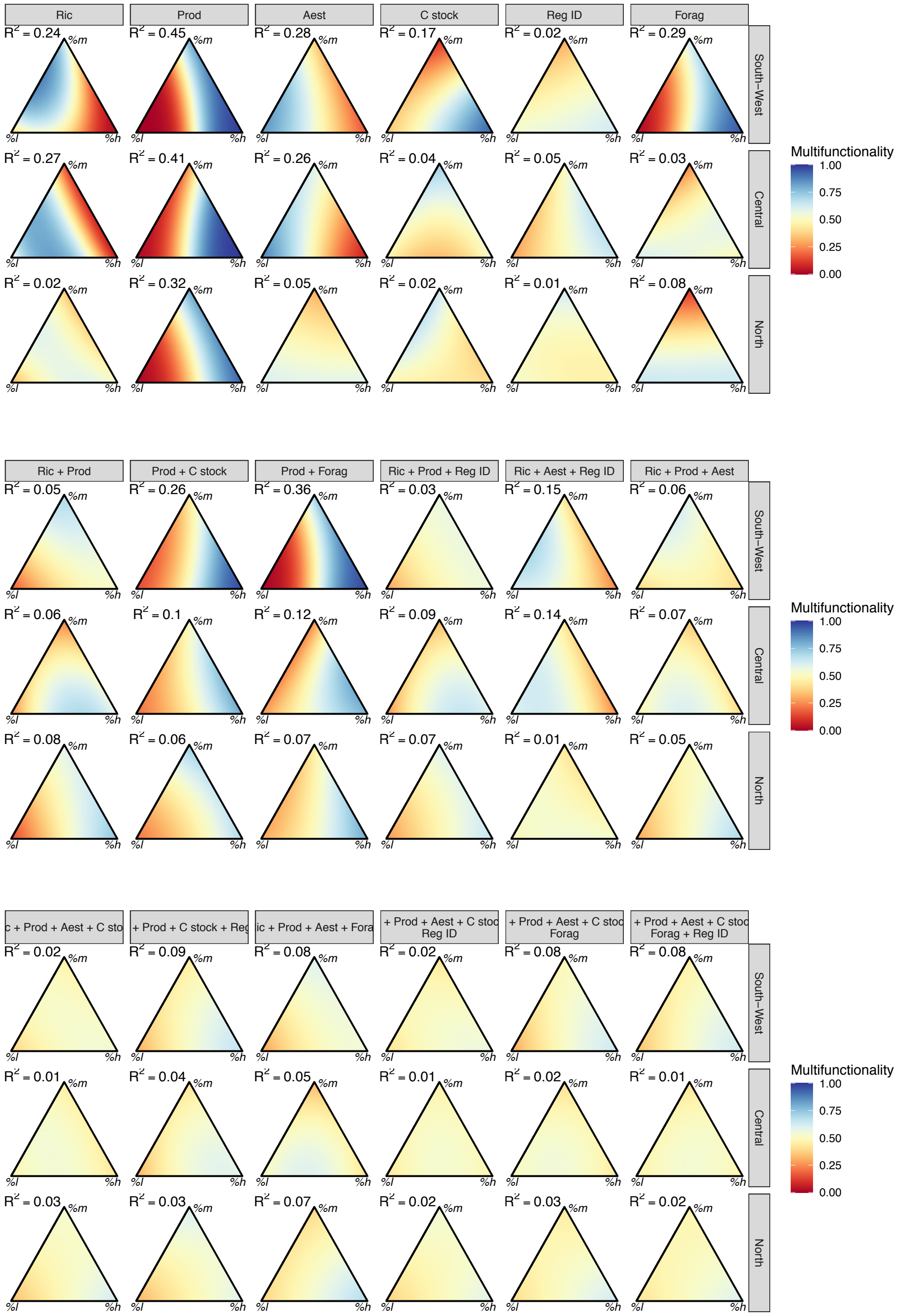

**Figure A3.** Estimated multifunctionality values for different stakeholder groups (column), depending on landscape composition (in proportions of low, medium and high-intensity sites) for the three regions (rows). Multifunctionality values are calculated by multiplying each service's threshold value (i.e., 1 if the service is above the median, 0 otherwise), weighted by its relative priority to the stakeholder group.

Note that the scale had been expanded (ranging from 0.3 to 0.8 instead of 0 to 1) compared to the other figures due to low multifunctionality ranges.

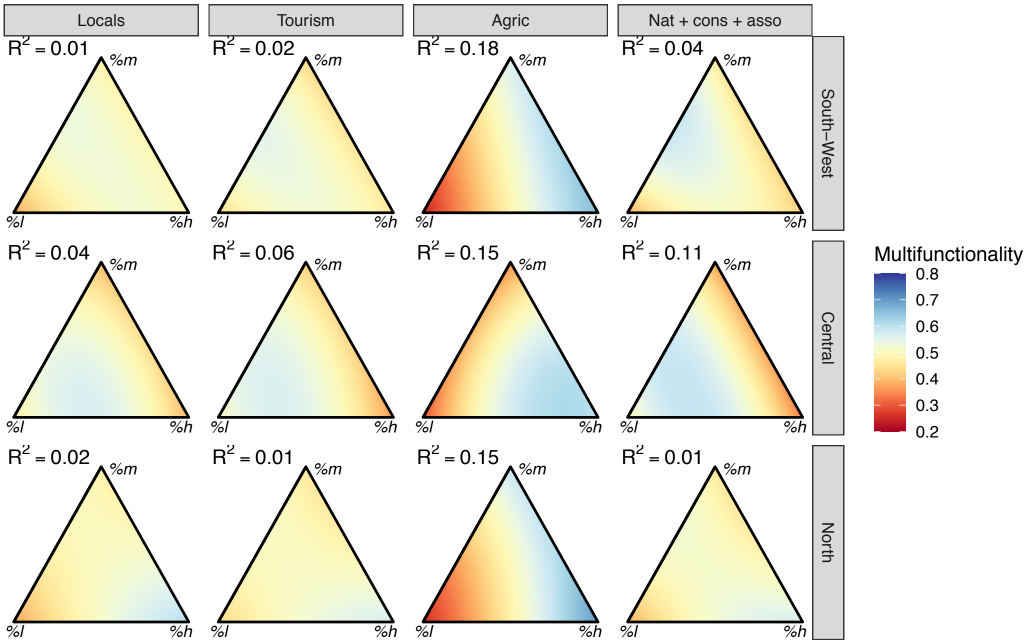
