## Supplementary methods for "Assessing the impact of grassland management on landscape multifunctionality"

### Appendix B: Supplementary methods

Neyret M., Fischer M., Allan E., Hölzel N., Klaus V. H., Kleinebecker T., Krauss J., Le Provost G., Peter. S., Schenk N., Simons N.K., van der Plas F., Binkenstein J., Börschig C., Jung K., Prati D., Schäfer D., Schäfer M., Schöning I., Schrumpf M., Tschapka M., Westphal C. & Manning P.

### Measurement of site-level indicators and aggregation to landscape level

The ‘biodiversity conservation’ service at the site-level was based on total plant species richness as plant alpha-diversity. Plant species richness has been shown to be a good proxy for diversity at multiple trophic levels at these sites (correlation of 0.67 and 0.68 between the whole ecosystem multidiversity index (Allan et al 2014) and the richness of asterids and rosids respectively, for instance (Manning et al., 2015)). We chose not to include other taxa to prevent co-linearity with the other service measures (see below). This indicator was calculated at the landscape level as gamma diversity.

**Figure B1 Correlations among potential service indicators**

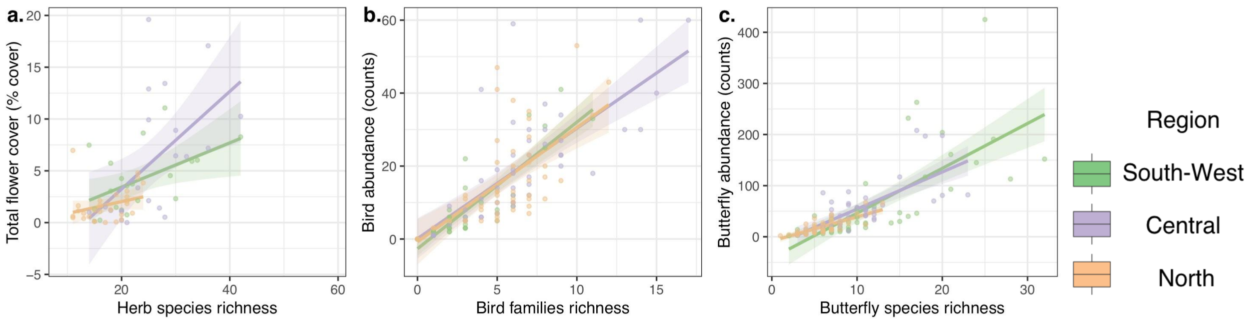


The fodder production service was calculated as total fodder protein production, a common agronomical indicator (Lee, 2018) that we calculated based on grassland aboveground biomass production and shoot protein content. Between mid-May and mid-June each year, aboveground biomass was harvested by clipping the vegetation 2 – 3 cm above ground in four randomly placed quadrats of 0.5 m × 0.5 m in each subplot. The plant biomass was dried at 80°C for 48 hours, weighed and summed over the four quadrats. Biomass was then averaged between 2008 and 2012. In order to convert this one-time biomass measurements into estimates of annual field productivity, we used the information on the number of cuts and the number of livestock units in a site to estimate the total biomass production used by farming activities, i.e. converted into fodder or consumed directly by livestock. Details of this estimation process can be found in Simons & Weisser (2017). We then multiplied this productivity by plant shoot protein levels, a common indicator of forage quality (Lee, 2018). Total nitrogen concentrations in ground samples of aboveground biomass were determined using an elemental auto-analyser (NA1500, CarloErba, Milan, Italy), and multiplied by 6.25 to obtain protein content (Lee, 2018). The landscape-scale protein production was then calculated as the sum of the production of all individual sites in the landscape.

Climate change mitigation was quantified as soil organic carbon stocks in the top 10 cm, as deeper stocks are unlikely to be affected strongly by management actions. We sampled composite samples for each site, prepared by mixing 14 mineral surface soil samples per site. Soil samples were taken along two 18 m transects in each site using a split tube auger, 40 cm long and 5 cm wide (Eijkelkamp, Giesbeek, The Netherlands). Composite samples were weighed, homogenized, air-dried and sieved (<2 mm). We then measured total carbon (TC) contents by dry combustion in a CN analyser "Vario Max" (Elementar Analysensysteme GmbH, Hanau, Germany) on ground subsamples. We determined inorganic carbon (IC) contents after combustion of organic carbon in a muffle furnace (450°C for 16 h). We then calculated the soil organic carbon (SOC) content as the difference between TC and IC, and the SOC concentration based on the weight of the dry fine-earth (105°C) and its volume. SOC concentration was then multiplied by soil bulk density to obtain site-level carbon stock values. The landscape-scale soil carbon stock was calculated as the sum of the soil carbon stock of all individual sites.

The aesthetic value measure integrated flower cover, the number of bird families and abundance of butterflies. The choice of these indicators was led by studies showing people's preference for bird richness over abundance (Cox & Gaston, 2015), including song diversity (Hedblom et al., 2014); and for flower-rich landscapes (Graves et al., 2017). Flowering units were counted between May and September 2009 for all flowering plant species (excluding grasses and sedges) on transects along the four edges of each site, in a total area of 600m$^2$. For abundant species, the number of flowering units was extrapolated to the whole site from a smaller area of 112 m$^2$. The total flower cover was calculated at the site scale as the sum of the individual flower cover of all plant species (see Binkenstein et al. (2013) for details). Butterfly and day-active moths (hereafter termed as Lepidoptera) abundance was measured in 2008 and averaged among sites within each landscape (Börschig et al., 2013). We conducted surveys of Lepidoptera from early May to mid-August. We sampled Lepidoptera during 3 surveys, each along one fixed 300m transect of 30min in each site. Each transect was divided in 50m sections of 5min intervals and we recorded all Lepidoptera within a 5 m corridor. Birds were surveyed by standardized audio-visual point-counts and all birds exhibiting territorial displays (singing and calling) were recorded. We used fixed-radius point counts and recorded all males of each bird species during a five-minute interval per site. Each site was visited five times between 15 March and 15 June each year. The data was then aggregated by family. Landscape-scale bird richness was calculated as the total number of bird families found in the landscape (i.e. in at least one site and one year) between 2009 and 2012. These three indicators were then scaled and averaged to estimate landscape-scale aesthetic value. Richness and abundance were usually highly correlated for the three groups (correlation of 0.75 for butterflies, 0.72 for birds, and 0.52 for plants), and the number of families for birds was highly correlated (0.96) to species richness; thus, other selections of indicators would have led to similar results (Fig. B1).

The regional identity service was estimated from the richness of culturally important birds and the cover of culturally important plants. The list of culturally important bird species was obtained from an online survey among 57 German respondents who were asked to say whether each species observed on the sites (see above) was unimportant (0 points), slightly important (1 point) or very important (5 points) species, based on whether the species "were part of German cultural identity or that of the study regions, e.g. by regularly appearing in German folklore, iconography, or popular entertainment". We then retained the 25% species with highest average scores (Table B1), and used the survey data described above to calculate species richness. The list of culturally important plant species also followed the same definition ("part of German cultural identity or that of the study regions, e.g. by regularly appearing in German folklore, iconography, or popular entertainment") but was created by two German experts with botanical knowledge of the species in the study regions (Table B2). We then used the survey data described above to calculate total cover of the species they listed.

Finally, the foraging service was calculated as the total cover of edible plants. The list of edible plants was obtained by the same botanical experts as the culturally important species and is presented in Table B3.

Table B1. List of culturally important birds

| | | |
|---|---|---|
| *Alauda arvensis* | *Falco tinnunculus* | *Milvus milvus* |
| *Buteo buteo* | *Fringilla coelebs* | *Parus major* |
| *Ciconia ciconia* | *Garrulus glandarius* | *Pica pica* |
| *Corvus corax* | *Grus grus* | *Sturnus vulgaris* |
| *Cuculus canorus* | *Hirundo rustica* | *Troglodytes troglodytes* |
| *Cyanistes caeruleus* | | *Turdus merula* |

Table B2. List of culturally important plants

| | | | |
|---|---|---|---|
| *Acer sp.* | *Colchicum autumnale* | *Prunus avium* | *Trifolium pratense* |
| *Bellis perennis* | *Equisetum arvense* | *Pseudolysimachion spicatum* | *Trifolium repens* |
| *Betula pendula* | *Fraxinus excelsior* | *Pulsatilla vulgaris* | *Urtica dioica* |
| *Briza media* | *Gentiana verna* | *Quercus robur* | *Veronica arvensis* |
| *Campanula glomerata* | *Gentianella germanica* | *Rosa canina aggr.* | *Veronica chamaedrys* |
| *Campanula patula* | *Geum urbanum* | *Taraxacum sp* | *Veronica filiformis* |
| *Campanula persicifolia* | *Juniperus communis* | *Thymus pulegioides* | *Veronica hederifolia* |

| | | | |
|---|---|---|---|
| *Campanula rapunculoides* | *Leucanthemum vulgare aggr.* | *Tilia sp.* | *Veronica officinalis* |
| *Campanula rotundifolia* | *Myosotis arvensis* | *Trifolium alpestre* | *Veronica persica* |
| *Campanula sp.* | *Myosotis discolor* | *Trifolium arvense* | *Veronica serpyllifolia* |
| *Carlina acaulis* | *Myosotis ramosissima* | *Trifolium campestre* | *Veronica teucrium* |
| *Carpinus betulus* | *Origanum vulgare* | *Trifolium dubium* | *Viola arvensis* |
| *Centaurium erythraea* | *Phragmites australis* | *Trifolium medium* | *Viola hirta* |
| *Cichorium intybus* | *Primula elatior veris agg* | *Trifolium montanum* | *Viola sp.* |

Table B3. List of edible plants

| | | | |
|---|---|---|---|
| *Achillea millefolium aggr.* | *Carum carvi* | *Lamium album* | *Rumex acetosella* |
| *Aegopodium podagraria* | *Chenopodium album* | *Lamium maculatum* | *Rumex thyrsiflorus* |
| *Allium cf oleraceum* | *Cichorium intybus* | *Leontodon hispidus* | *Salvia pratensis* |
| *Angelica sylvestris* | *Cirsium oleraceum* | *Lepidium campestre* | *Sanguisorba minor* |
| *Anthriscus sylvestris* | *Crataegus sp.* | *Origanum vulgare* | *Silene dioica* |
| *Aphanes arvensis* | *Daucus carota* | *Pastinaca sativa* | *Silene vulgaris* |
| *Arctium lappa* | *Elymus repens* | *Phyteuma orbiculare* | *Sinapis alba* |
| *Arctium minus* | *Erigeron acris* | *Phyteuma spicatum* | *Sinapis arvensis* |
| *Arctium sp.* | *Fallopia convolvulus* | *Prunus avium* | *Sisymbrium officinale* |
| *Arctium tomentosum* | *Filipendula ulmaria* | *Prunus sp.* | *Stellaria media* |
| *Astragalus glycyphyllos* | *Fragaria vesca* | *Prunus spinosa* | *Symphytum officinale* |
| *Bellis perennis* | *Fragaria viridis* | *Ranunculus ficaria* | *Taraxacum sp.* |
| *Bistorta officinalis* | *Glechoma hederacea* | *Rosa canina aggr.* | *Thlaspi arvense* |
| *Brassica rapa* | *Heracleum sphondylium* | *Rubus caesius* | *Thymus pulegioides* |
| *Bunium bulbocastanum* | *Hypericum perforatum* | *Rubus sp.* | *Tragopogon pratensis* |
| *Cardamine pratensis* | *Juniperus communis* | *Rumex acetosa* | |

### Topographical Wetness Index calculation

We calculated the Topographic Wetness Index (TWI) of each site, defined as $\ln(a/\tan B)$ where $a$ is the specific catchment area (cumulative upslope area which drains through a Digital Elevation Model (DEM, http://www.bkg.bund.de) cell, divided by per unit contour length) and $\tan B$ is the slope gradient in radians calculated over a local region surrounding the cell of interest (Gessler et al. 1995; Sørensen et al. 2006). TWI therefore combines both upslope contributing area (determining the amount of water received from upslope areas) and slope (determining the loss of water from the site to downslope areas). TWI was calculated from raster DEM data with a cell size of 25 m for all sites, using ArcGIS tools (flow direction and flow accumulation tools of the hydrology toolset and raster calculator). The TWI measure used was the average value for a 4 × 4 window centred on the site, i.e. 16 DEM cells corresponding to an area of 100 m ×100 m. See Le Provost et al. 2021 for further details.
