## Supplementary material for "Assessing the impact of grassland management on landscape multifunctionality": Sensitivity analyses

### Appendix C: Sensitivity analyses

Neyret M., Fischer M., Allan E., Hölzel N., Klaus V. H., Kleinebecker T., Krauss J., Le Provost G., Peter. S., Schenk N., Simons N.K., van der Plas F., Binkenstein J., Börschig C., Jung K., Prati D., Schäfer D., Schäfer M., Schöning I., Schrumpf M., Tschapka M., Westphal C. & Manning P.

We explored how multiple choices in the simulations and calculation of multifunctionality could affect the results of our analyses. In the first part of this appendix, we first list all the different sensitivities that we identified and detail how their implementation modified the methodology detailed in the main text. In the second part, we consider in turn each of the main results presented in the paper and describe the potential variations highlighted by the analyses. We describe only analyses departing from the main analysis by one parameter – further combinations can be explored in the online tool.

### 1. Methods

The sensitivities we identified were the following:

a. Analysis on non-environmentally corrected data.

The main analysis was conducted on service indicator values that were first corrected at the site level by environmental covariates (see Methods). In these sensitivity analyses we also conduct the analyses using raw data.

b. Classification into classes of low, medium and high land-use intensity.

In the main analyses, the sites are classified in the three intensity categories based on 33% quantiles of the intensity values (e.g. lowest, medium and highest thirds) within the regions. In these sensitivity analyses we remove the "intermediate", potentially confounding sites by using sites within the lowest, medium and highest 20% quantiles (e.g. lowest, medium and highest fifths) of the intensities within each region. This reduced the number of sites available during the landscape simulations, and to avoid running the analysis on identical landscapes we adapted the analyses by lowering the number of landscape replicates for each combination (*).

c. Number of sites per landscape.

In the main analysis, each landscape was composed of 10 sites. We also run similar analyses with 7 and 13 sites per landscapes. This also affected the number of possible landscape combinations, and we consequently adapted the number of landscape replicates for each combination (*).

d. Calculation of landscape-scale service indicator values.

In the main analysis, landscape-scale service indicator values consisted either of gamma diversity (plant and bird diversity) or of the sum of the services provided by all sites in the landscape (other services). We also considered a situation in which high level of each service is expected in only part of the landscape, and calculated the landscape indicator value as the maximum of the indicator in all sites of the landscape.

e. Calculation of landscape-scale multifunctionality

In the main analysis, landscape scale multifunctionality was calculated as the number of services above the median or as 1 if all services were over 25%, 0 otherwise. We also calculated multifunctionality as the number of services above the $40^{th}$ or $60^{th}$ quantiles of the distribution of the values in all landscapes (measured as $95^{th}$ quantile to avoid outliers).

We also measured multifunctionality as the average of the (scaled) values for all considered services.

(*) In the main analysis, there were 66 possible landscape compositions (from 0 to 10 sites of each intensity), and 15 random landscape replicates per composition, hence 990 different landscapes. Changing the number of sites per landscapes changed the number of possible combinations (36 possible combinations for 7 sites, 105 for 13 sites) and to keep the total number of simulated landscapes approximately similar, we used 1000/(number of combinations) landscape replicates per combination (i.e. 10 for 13 sites, 28 for 7 sites).

When using only the 20% lowest, medium and highest intensity sites decreased the size of the regional pool from which to build landscapes. In that case we used only 7 sites per landscape and the extreme compositions (e.g. 100% of one intensity) were represented by slightly less landscape replicates.

### 2. Results

a. Site-level correlations among indicators and variation of service indicator values with land-use intensity and.

When correcting for the environment, there were positive correlations between flower cover, butterfly abundance and plant species richness. These indicators were usually positively correlated with bird richness and negatively correlated with biomass production. Most of these relationships were similar when considering raw, non environmentally-corrected data (Table C1). As shown in Table C1, the variation of site-level service indicators with land-use intensity was not strongly affected by using raw data instead of environmentally corrected residuals, except that the response of some services (e.g. plant richness) to intensity was more linear (no apparent "threshold") when it was corrected for the environment.

b. Landscape-level correlations among ecosystem services.

Correlations among landscape-scale services was mostly similar when considering services calculated as the maximum value of each service (instead of the sum) in the landscape (Fig. C3), when changing the number of sites per landscape (Fig. C4, Fig. C5) or using data not corrected for the environment (Fig. C6).

c. Multifunctionality response to landscape composition

The following figures present the multifunctionality response to landscape composition as affected by the parameters of the model.
Calculating the landscape-scale services as the maximum instead of the sum (Fig. C8) did not change the direction of the response. It led to weaker responses of single services (top row) and marginally stronger variability when including multiple services.
Decreasing (Fig. C9) or increasing (Fig. C10) the number of sites per landscape slightly weakened (resp. strengthened) the response of single services to landscape composition, possibly because including more sites made for a higher chance to select sites with very high or low values, especially in the extreme compositions (e.g. 100% low intensity, 100% high intensity). When considering multiple services, including more sites did not change the response of multifunctionality.
For multifunctionality counted as the proportion of services above a given threshold, changing the threshold from the median to the 40$^{th}$ or 60$^{th}$ quantile of the distribution slightly switched the multifunctionality to higher or, respectively, lower values (Fig. C11 and Fig. C12) but the general form of the response was not affected.
The same was observed in the “compromise” scenario, when multifunctionality was calculated as 1 when all services were above a threshold, and 0 otherwise. Changing the threshold from the 25$^{th}$ quantile of the distribution to the 15$^{th}$ or 35$^{th}$ quantiles (Fig. C13 and Fig. C14) made it easier (respectively, more difficult) to provide all services at the required level, resulting in overall higher (resp. lower) multifunctionality values but without affecting the form of the response.
Calculating multifunctionality as the average of all services gave similar results as the main thresholding approach (Fig. C15).

d. Effect of other drivers on the responsivity of multifunctionality to landscape composition

Table C2 presents the result of the model with the responsivity (i.e. range) of multifunctionality over all landscape compositions as a response, and the ratio of the effect of land-use intensity and other environmental variables; the number of services included; or the service response variance as explanatory variables.
The relative effect of land-use intensity compared to other environmental variables was calculated as the ratio between the slope coefficient between individual services and land-use intensity over the maximum slope coefficient between the service and all other environmental covariates. It was significantly positive regardless of the model parameters, except when landscape-scale services were calculated based on the maximum of the sites.
The multifunctionality range decreased with the number of services considered in 20 out of the 22 scenarios considered, supporting our main conclusions. However, it did not vary with the number of services when multifunctionality was calculated as 1 if all the services were above the

15th percentile. This is because for low thresholds such as these, multifunctionality is expected to be high everywhere if too few services are considered.

Finally, the responsiveness of multifunctionality significantly decreased with the service response variance in 18 of the 22 considered scenarios. There was no positive relationship.

**Table C1 Site-level correlation among ecosystem services.** a. corrected for the environment, and b. raw service values. * P < 0.05, ** P < 0.01, *** P < 0.001.

| **a. Indicators corrected for the environment** | Biomass production | Plant nitrogen content | Flower cover (sqrt) | Butterfly abundance (sqrt) | Bird family richness | Plant richness | Soil C stock | Cover edible plants | *Juniperus* grasslands | Cover cultural plants |
|---|---|---|---|---|---|---|---|---|---|---|
| Plant nitrogen content | 0.16 | | | | | | | | | |
| Flower cover (sqrt) | 0.02 | -0.06 | | | | | | | | |
| Butterfly abundance (sqrt) | -0.48 *** | -0.16 | 0.17 * | | | | | | | |
| Bird family richness | -0.25 ** | -0.07 | -0.03 | 0.28 *** | | | | | | |
| Plant richness | -0.5 *** | -0.07 | 0.41 *** | 0.47 *** | 0.35 *** | | | | | |
| Soil C stock | 0.02 | 0.11 | -0.02 | -0.02 | 0.04 | -0.09 | | | | |
| Cover edible plants | 0.17 * | 0.08 | -0.04 | -0.16 | -0.01 | -0.15 | -0.12 | | | |
| *Juniperus* grasslands | -0.28 * | -0.01 | -0.04 | 0.31 ** | 0.29 * | 0.24 | 0.07 | -0.09 | | |
| Cover cultural plants | 0.28 *** | 0.01 | 0.16 * | -0.19 * | -0.03 | -0.14 | -0.17 * | 0.38 *** | -0.1 | |
| Richness cultural birds | -0.18 * | -0.05 | 0.02 | 0.23 ** | 0.79 *** | 0.27 *** | 0.05 | -0.09 | 0.29 *** | 0.04 |

| **b. Raw indicator values** | Biomass production | Plant nitrogen content | Flower cover (sqrt) | Butterfly abundance (sqrt) | Bird family richness | Plant richness | Soil C stock | Cover edible plants | *Juniperus* grasslands | Cover cultural plants |
|---|---|---|---|---|---|---|---|---|---|---|
| Plant nitrogen content | 0.19 * | | | | | | | | | |
| Flower cover (sqrt) | -0.11 | -0.12 | | | | | | | | |
| Butterfly abundance (sqrt) | -0.53 *** | -0.2 * | 0.36 *** | | | | | | | |
| Bird family richness | -0.29 *** | -0.11 | -0.03 | 0.26 ** | | | | | | |
| Plant richness | -0.53 *** | -0.17 * | 0.62 *** | 0.6 *** | 0.31 *** | | | | | |
| Soil C stock | 0.14 | 0.15 | -0.31 *** | -0.3 *** | -0.04 | -0.43 *** | | | | |
| Cover edible plants | 0.21 ** | 0.07 | -0.19 * | -0.25 ** | -0.01 | -0.27 ** | 0.12 | | | |
| *Juniperus* grasslands | -0.24 ** | -0.07 | 0.04 | 0.38 *** | 0.22 ** | 0.24 ** | -0.09 | -0.07 | | |
| Cover cultural plants | 0.24 ** | 0.02 | 0.25 ** | -0.05 | -0.08 | 0.01 | -0.3 *** | 0.22 ** | -0.04 | |
| Richness cultural birds | -0.24 ** | -0.12 | 0.05 | 0.27 *** | 0.82 *** | 0.3 *** | -0.13 | -0.1 | 0.26 ** | 0.03 |

**Table C2 Variation in ecosystem services and their component indicators with the land-use intensity class in each region (mean ± sd)**. Indicators were not corrected for the environment (see Methods). Different letters indicate differences significant at 5%.

| **Ecosystem service** or *service indicator* | Land-use intensity | South-West | Central | North |
|---|---|---|---|---|
| **Livestock production** | low | 52.07 +/- 18.45 [a] | 50.91 +/- 13.94 [a] | 105.9 +/- 15.08 [a] |
| | medium | 167.19 +/- 19.02 [b] | 98.04 +/- 14.37 [a] | 141.2 +/- 15.54 [ab] |
| | high | 189.16 +/- 18.45 [b] | 163.26 +/- 13.94 [b] | 169.04 +/- 15.08 [b] |
| *Biomass production* | low | 23.66 +/- 6.92 [a] | 25.82 +/- 5.99 [a] | 48 +/- 5.47 [a] |
| | medium | 73.01 +/- 7.13 [b] | 46.36 +/- 6.18 [a] | 64.78 +/- 5.64 [ab] |
| | high | 88.68 +/- 6.92 [b] | 78.39 +/- 5.99 [b] | 72.74 +/- 5.47 [b] |
| *N content* | low | 2.12 +/- 0.08 [a] | 1.95 +/- 0.07 [a] | 2.24 +/- 0.13 [a] |
| | medium | 2.18 +/- 0.08 [a] | 2.1 +/- 0.07 [a] | 2.18 +/- 0.14 [a] |
| | high | 2.12 +/- 0.08 [a] | 2.08 +/- 0.07 [a] | 2.27 +/- 0.13 [a] |
| **Aesthetic** | low | 0.49 +/- 0.03 [b] | 0.51 +/- 0.03 [b] | 0.26 +/- 0.02 [a] |
| | medium | 0.29 +/- 0.03 [a] | 0.43 +/- 0.04 [ab] | 0.28 +/- 0.02 [a] |
| | high | 0.3 +/- 0.03 [a] | 0.32 +/- 0.03 [a] | 0.25 +/- 0.02 [a] |
| *Flower cover (sqrt)* | low | 2.56 +/- 0.21 [b] | 2.53 +/- 0.26 [b] | 1 +/- 0.16 [a] |
| | medium | 1.68 +/- 0.21 [a] | 2.13 +/- 0.26 [ab] | 1.36 +/- 0.17 [a] |
| | high | 2.28 +/- 0.21 [ab] | 1.63 +/- 0.26 [a] | 1.18 +/- 0.16 [a] |
| *Butterfly abundance (sqrt)* | low | 10.32 +/- 0.81 [b] | 8.47 +/- 0.81 [b] | 4.32 +/- 0.43 [a] |
| | medium | 5.49 +/- 0.84 [a] | 6.84 +/- 0.84 [ab] | 4.24 +/- 0.44 [a] |
| | high | 4.12 +/- 0.81 [a] | 5.14 +/- 0.81 [a] | 4.2 +/- 0.43 [a] |
| *Bird family richness* | low | 4.59 +/- 0.56 [b] | 6.53 +/- 0.75 [a] | 5.06 +/- 0.51 [a] |
| | medium | 2.88 +/- 0.58 [ab] | 6.25 +/- 0.77 [a] | 4.56 +/- 0.52 [a] |
| | high | 2.47 +/- 0.56 [a] | 4.65 +/- 0.75 [a] | 4.06 +/- 0.51 [a] |
| **Conservation** *(plant richness)* | low | 54.88 +/- 2.01 [b] | 59.88 +/- 2.87 [b] | 27 +/- 1.27 [a] |
| | medium | 36.5 +/- 2.07 [a] | 44.69 +/- 2.96 [a] | 31 +/- 1.31 [a] |
| | high | 37.35 +/- 2.01 [a] | 38 +/- 2.87 [a] | 29.24 +/- 1.27 [a] |
| **C stock** | low | 3.75 +/- 0.18 [a] | 3.1 +/- 0.2 [a] | 6.49 +/- 0.79 [a] |
| | medium | 4.36 +/- 0.19 [ab] | 3.57 +/- 0.21 [a] | 4.79 +/- 0.82 [a] |
| | high | 4.57 +/- 0.18 [b] | 3.5 +/- 0.2 [a] | 5.32 +/- 0.79 [a] |
| **Foraging** *(Cover edible plants)* | low | 60.64 +/- 12.13 [a] | 87.25 +/- 12.28 [a] | 148.6 +/- 23.66 [a] |
| | medium | 105.07 +/- 12.51 [b] | 90.03 +/- 12.66 [a] | 81.67 +/- 24.39 [a] |
| | high | 122.56 +/- 12.13 [b] | 95.84 +/- 12.28 [a] | 130.83 +/- 23.66 [a] |
| **Regional ID** | low | 0.33 +/- 0.04 [b] | 0.23 +/- 0.03 [a] | 0.19 +/- 0.03 [a] |
| | medium | 0.19 +/- 0.04 [a] | 0.26 +/- 0.03 [a] | 0.22 +/- 0.03 [a] |
| | high | 0.22 +/- 0.04 [ab] | 0.28 +/- 0.03 [a] | 0.18 +/- 0.03 [a] |
| *Juniperus grasslands* | low | 0.24 +/- 0.07 [a] | 0 +/- 0 [a] | 0 +/- 0 [a] |
| | medium | 0.06 +/- 0.07 [a] | 0 +/- 0 [a] | 0 +/- 0 [a] |
| | high | 0 +/- 0.07 [a] | 0 +/- 0 [a] | 0 +/- 0 [a] |
| *Cover cultural plants* | low | 114.42 +/- 15.43 [a] | 77.58 +/- 14.72 [a] | 69.17 +/- 20.6 [a] |
| | medium | 93.27 +/- 15.91 [a] | 98.6 +/- 15.17 [a] | 113.05 +/- 21.23 [a] |
| | high | 145.01 +/- 15.43 [a] | 126.26 +/- 14.72 [a] | 100.92 +/- 20.6 [a] |
| *Richness cultural birds* | low | 2.29 +/- 0.32 [b] | 2.65 +/- 0.41 [a] | 2 +/- 0.26 [a] |
| | medium | 1.12 +/- 0.33 [a] | 2.69 +/- 0.42 [a] | 1.75 +/- 0.27 [a] |
| | high | 1.06 +/- 0.32 [a] | 2.41 +/- 0.41 [a] | 1.47 +/- 0.26 [a] |

**Figure C3 Trade-offs between landscape-scale ecosystem service measures**. This figure differs from Fig. 3 as the landscape-scale services were calculated based on the maximum (instead of sum) of the services provided by all sites in the landscape.

The colour and size of the circles denote the strength of the correlation between pairs of variables, within each region. Crosses indicate no significant correlations at 5% (Holm correction for multiple testing).

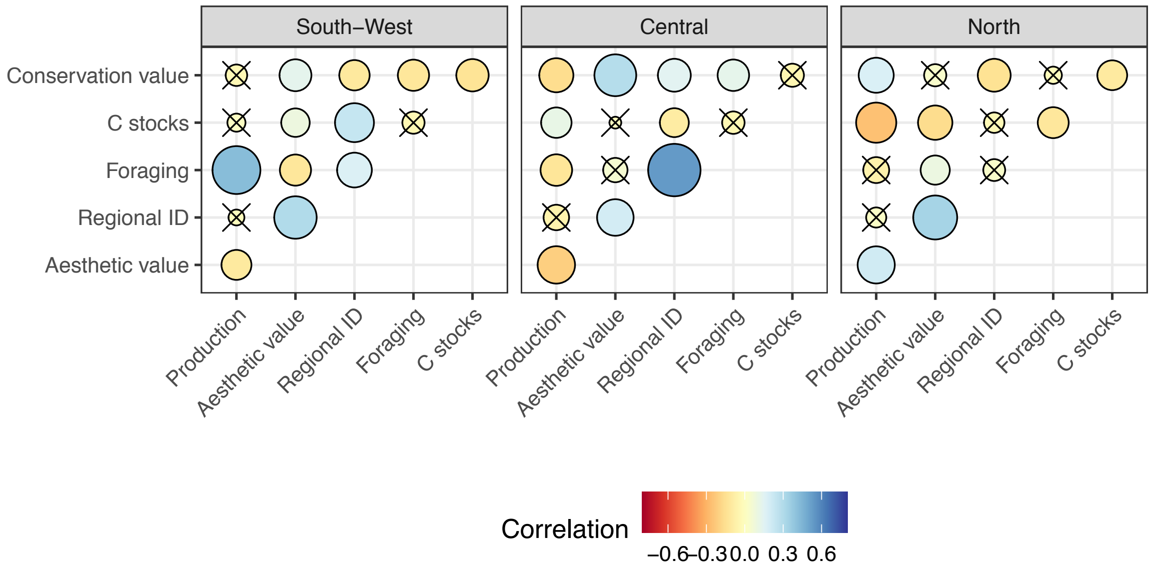


**Figure C4 Trade-offs between landscape-scale ecosystem service measures.** This figure differs from Fig. 3 as the landscapes included 7 (instead of 10) sites.

The colour and size of the circles denote the strength of the correlation between pairs of variables, within each region. Crosses indicate no significant correlations at 5% (Holm correction for multiple testing).

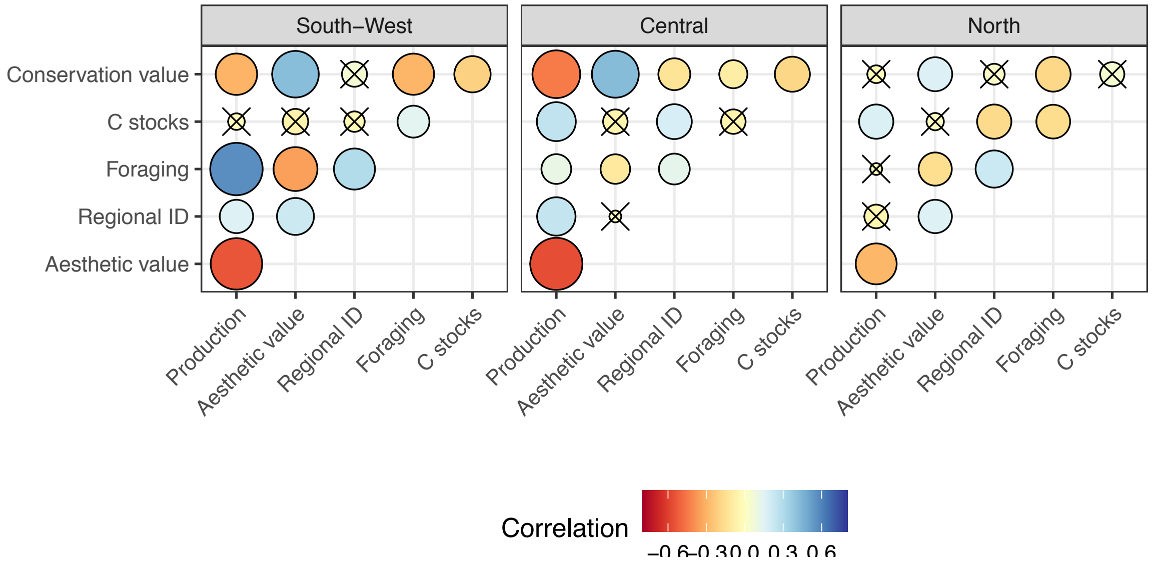

**Figure C5 Trade-offs between landscape-scale ecosystem service measures.** This figure differs from Fig. 3 as the landscapes included 13 (instead of 10) sites.

The colour and size of the circles denote the strength of the correlation between pairs of variables, within each region. Crosses indicate no significant correlations at 5% (Holm correction for multiple testing).

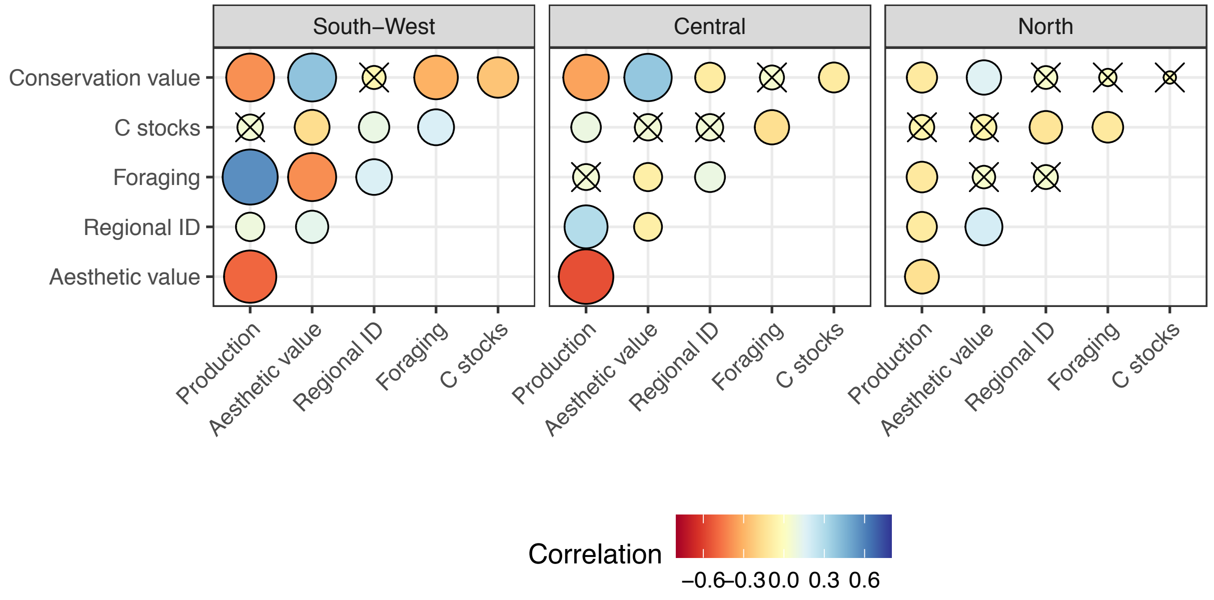


**Figure C6 Trade-offs between landscape-scale ecosystem service measures.** This figure differs from Fig. 3 as the ecosystem service indicators were not corrected for the environment before analysis.

The colour and size of the circles denote the strength of the correlation between pairs of variables, within each region. Crosses indicate no significant correlations at 5% (Holm correction for multiple testing).

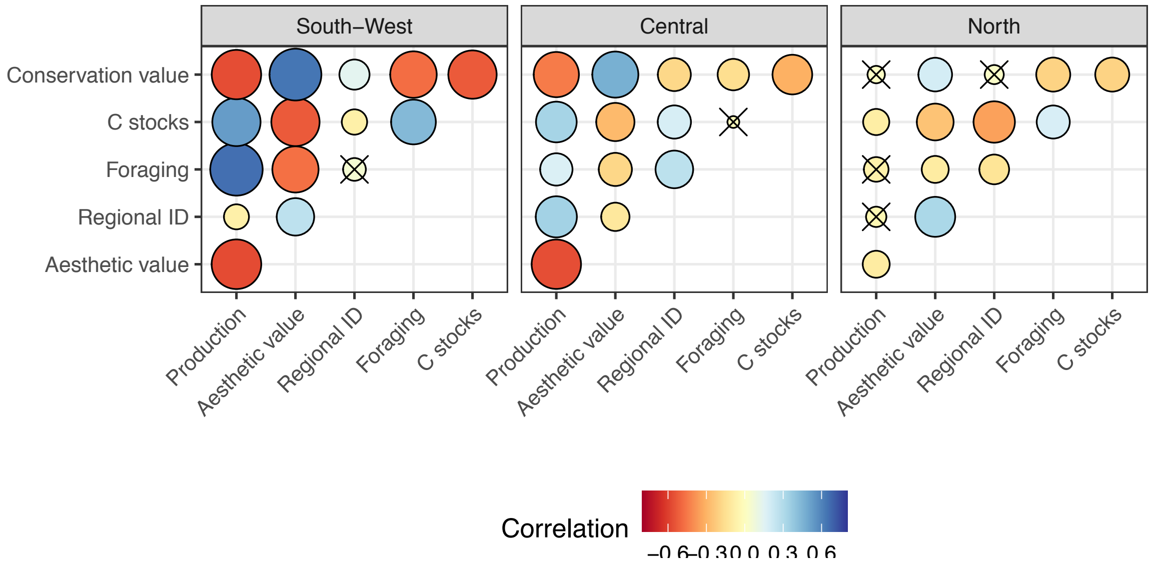

**Figure C7 Estimated multifunctionality values depending on landscape composition (in proportions of low, medium and high-intensity sites).** This figure shows all the service combinations for the 'compromise' approach, partly shown in Fig. 5.

For single ecosystem services (top row), the value presented corresponds to the probability of the given service to be above the median. For combinations of multiple services (middle and bottom rows), multifunctionality is the expected proportion of services above the median. Blue indicates higher multifunctionality values, orange lower.

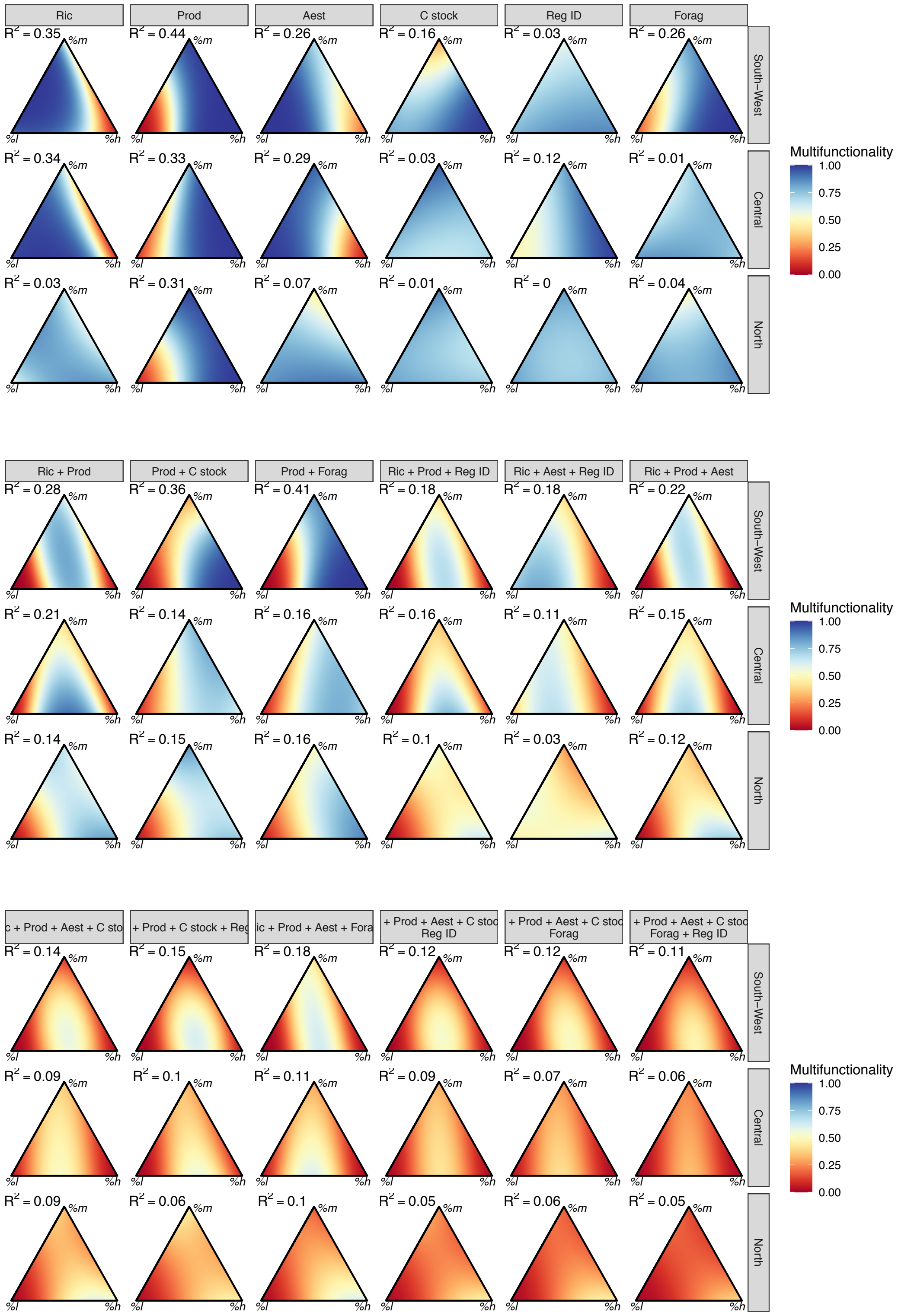

**Figure C8 Estimated multifunctionality values depending on landscape composition (in proportions of low, medium and high-intensity sites).** This figure differs from Fig. 4 in that landscape-scale ecosystem service values were calculated as the maximum, not the sum, of site-level ecosystem services.

For single ecosystem services (top row), the value presented corresponds to the probability of the given service to be above the median. For combinations of multiple services (middle and bottom rows), multifunctionality is the expected proportion of services above the median. Blue indicates higher multifunctionality values, orange lower.

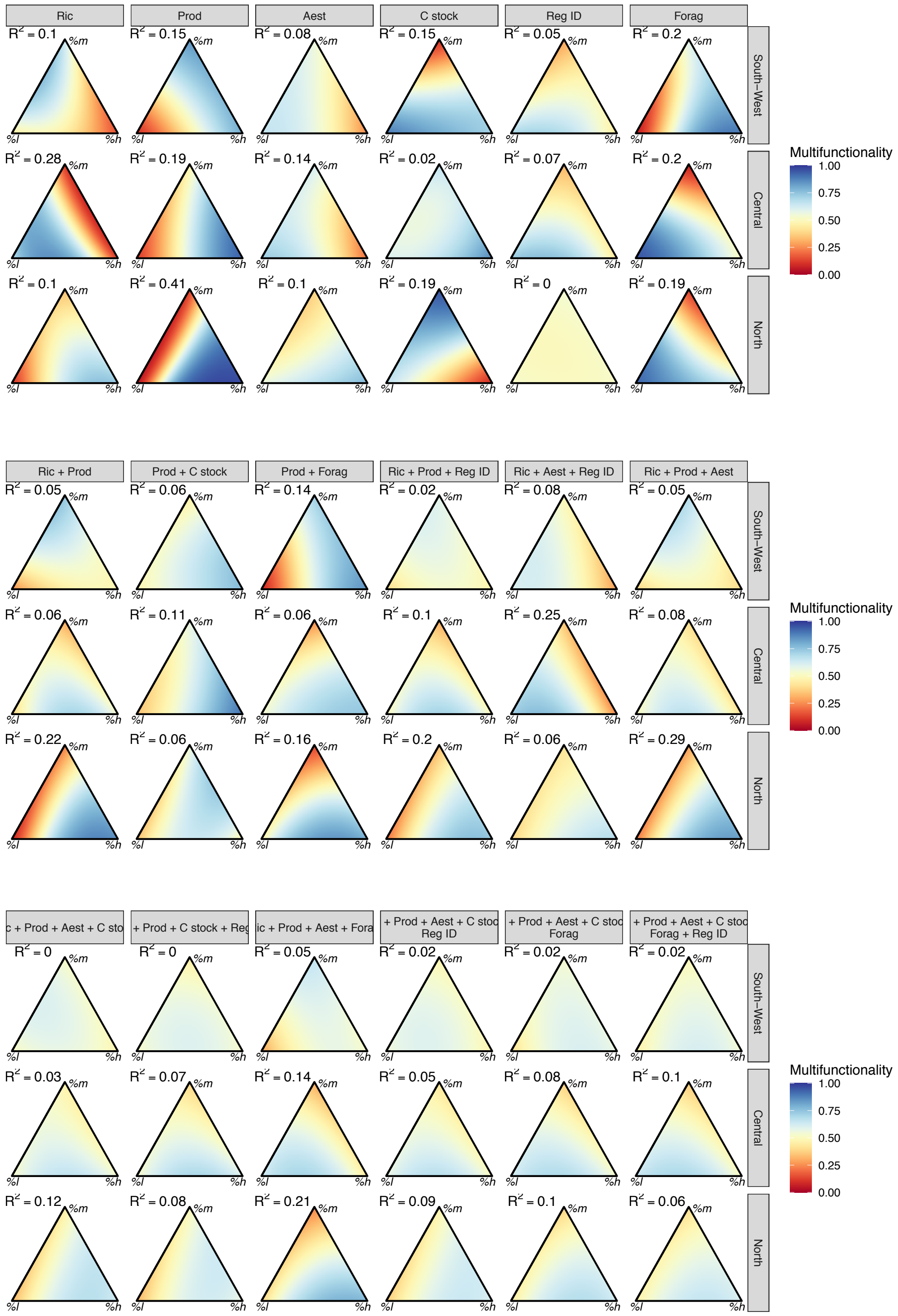

**Figure C9 Estimated multifunctionality values depending on landscape composition (in proportions of low, medium and high-intensity sites).** This figure differs from Fig. 4 in that landscapes were composed of 7 sites, instead of 10.

For single ecosystem services (top row), the value presented corresponds to the probability of the given service to be above the median. For combinations of multiple services (middle and bottom rows), multifunctionality is the expected proportion of services above the median. Blue indicates higher multifunctionality values, orange lower.

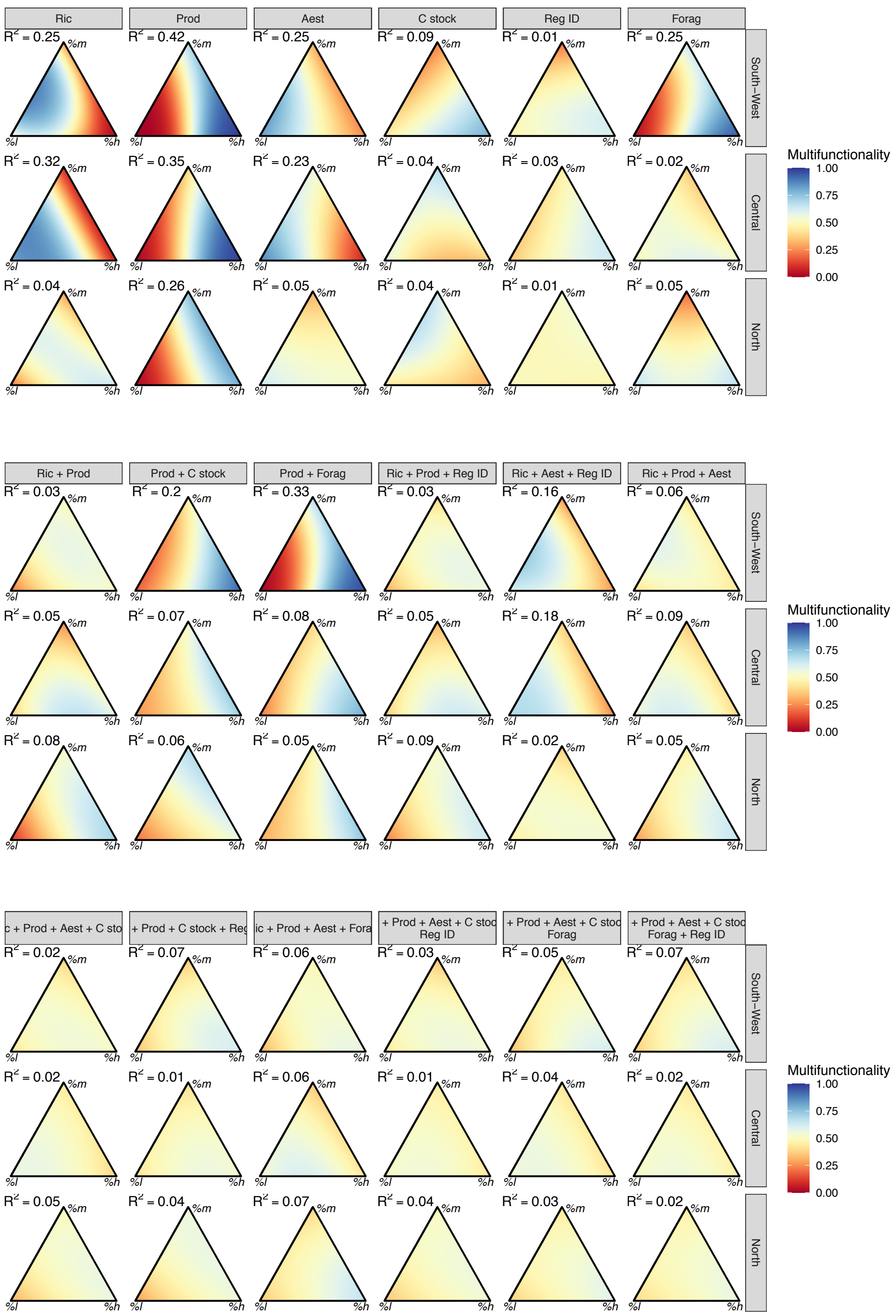

**Figure C10 Estimated multifunctionality values depending on landscape composition (in proportions of low, medium and high-intensity sites).** This figure differs from Fig. 4 in that landscapes were composed of 13 sites, instead of 10.

For single ecosystem services (top row), the value presented corresponds to the probability of the given service to be above the median. For combinations of multiple services (middle and bottom rows), multifunctionality is the expected proportion of services above the median. Blue indicates higher multifunctionality values, orange lower.

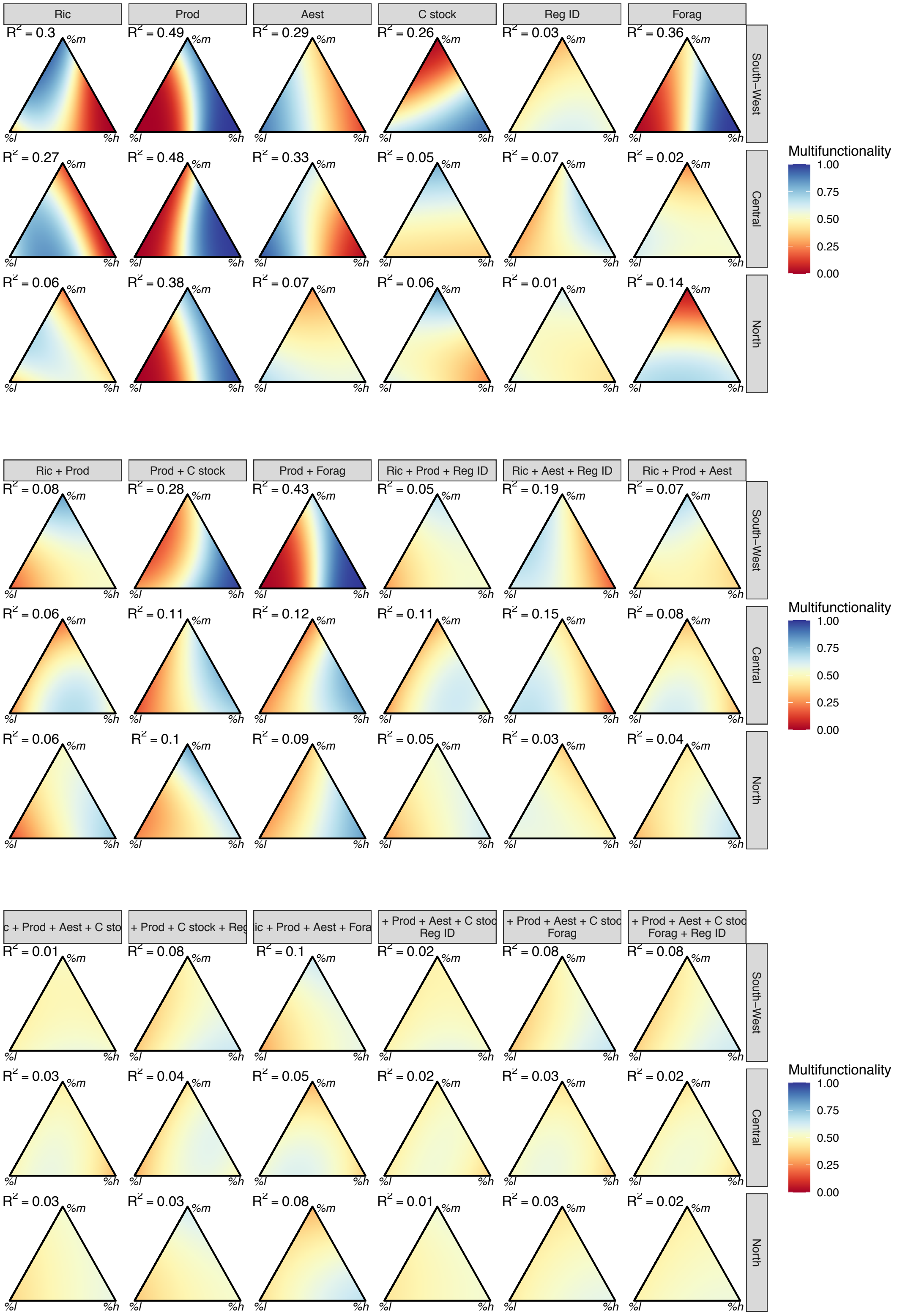

**Figure C11 Estimated multifunctionality values depending on landscape composition (in proportions of low, medium and high-intensity sites).** This figure differs from Fig. 4 in that the threshold was set to the 40th percentile instead of the median.

For single ecosystem services (top row), the value presented corresponds to the probability of the given service to be above the threshold. For combinations of multiple services (middle and bottom rows), multifunctionality is the expected proportion of services above the threshold. Blue indicates higher multifunctionality values, orange lower.

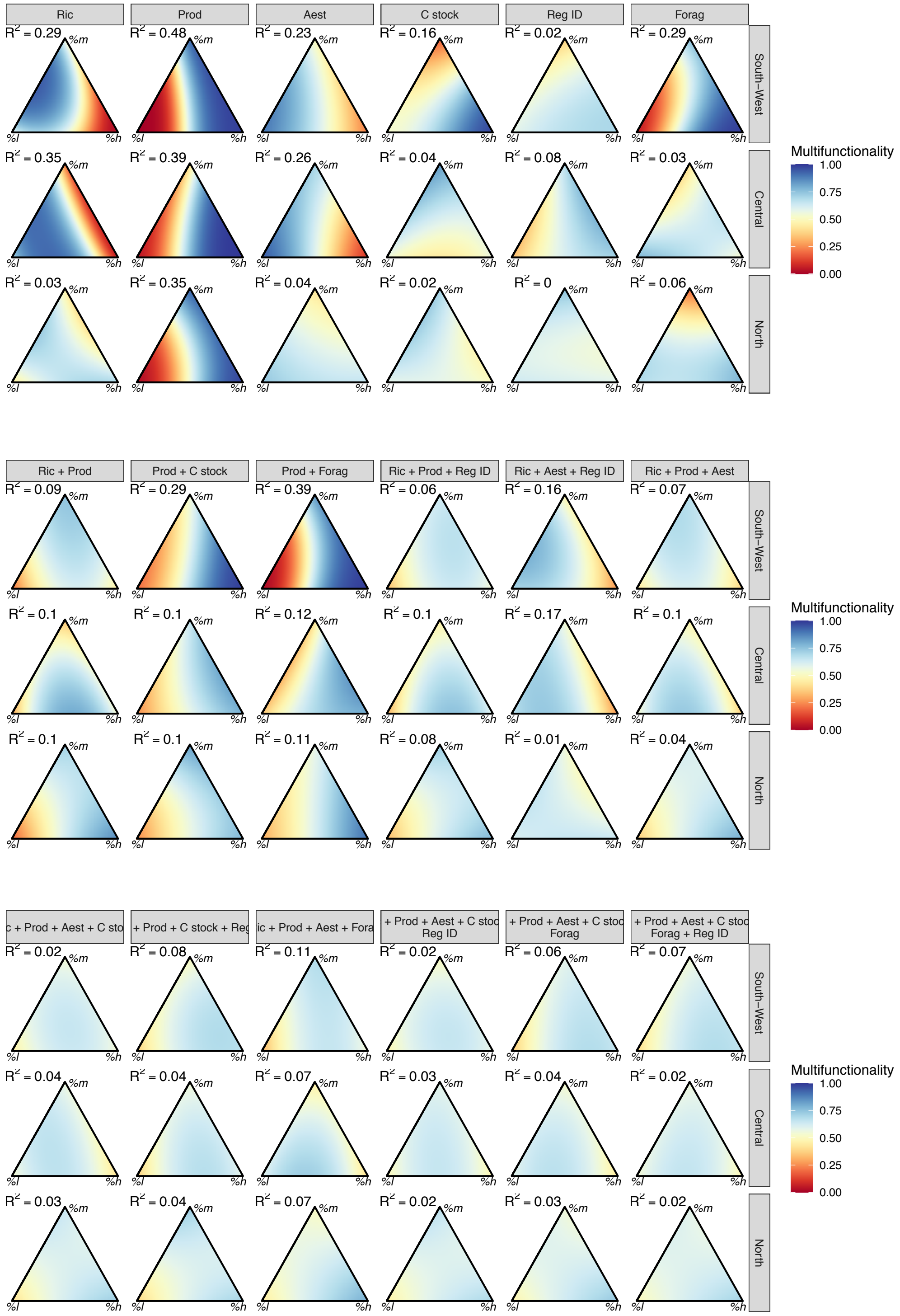

**Figure C12 Estimated multifunctionality values depending on landscape composition (in proportions of low, medium and high-intensity sites).** This figure differs from Fig. 4 in that the threshold was set to the 60th percentile instead of the median.

For single ecosystem services (top row), the value presented corresponds to the probability of the given service to be above the threshold. For combinations of multiple services (middle and bottom rows), multifunctionality is the expected proportion of services above the threshold. Blue indicates higher multifunctionality values, orange lower.

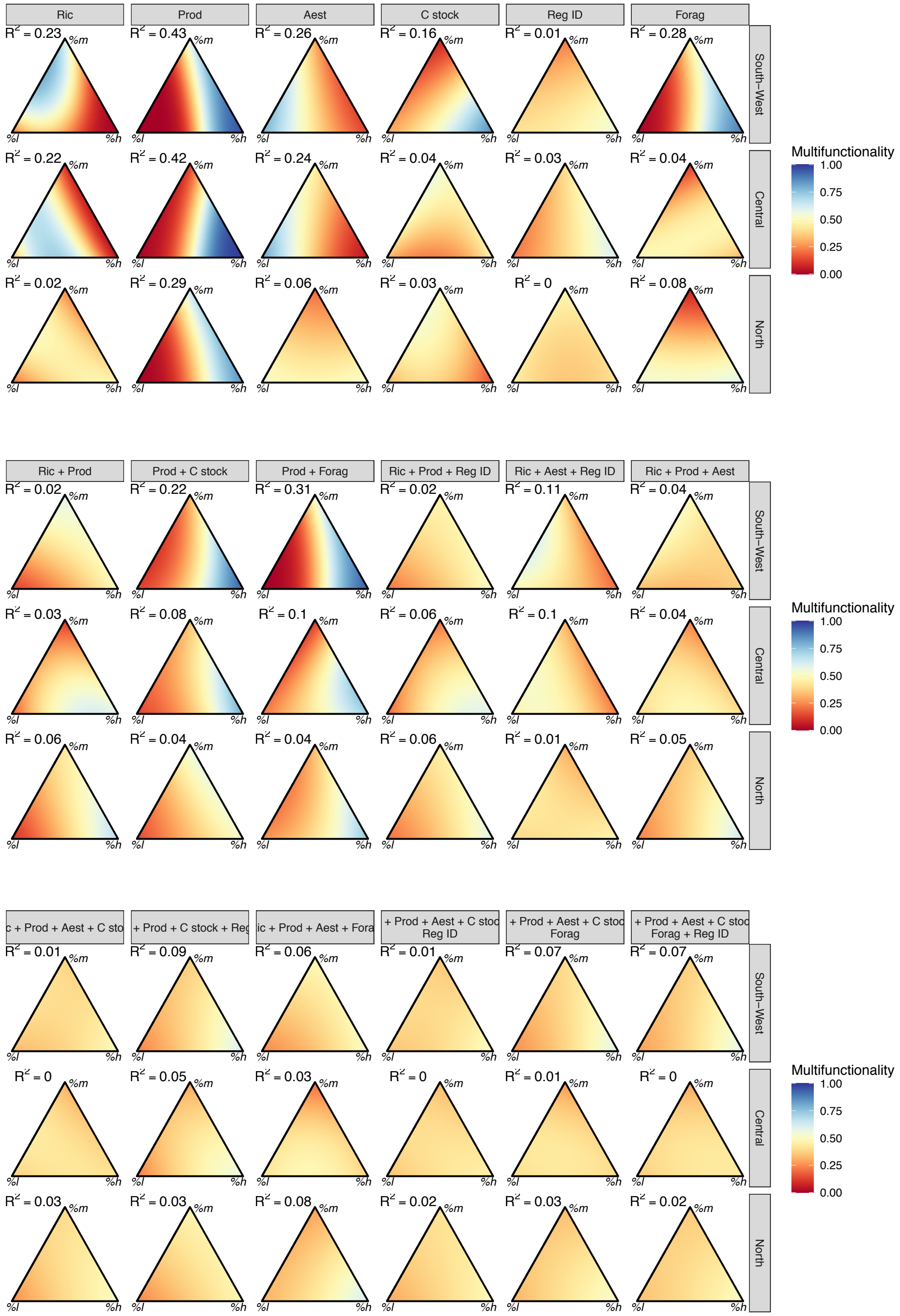

**Figure C13 Estimated multifunctionality values depending on landscape composition (in proportions of low, medium and high-intensity sites).** This figure differs from Fig. 5 in that the multifunctionality was calculated as 1 if all the services were above a 15th percentile threshold, and 0 otherwise (instead of a 25th percentile threshold).

The value presented corresponds to the probability that all given services are above the threshold. Blue indicates higher multifunctionality values, orange lower.

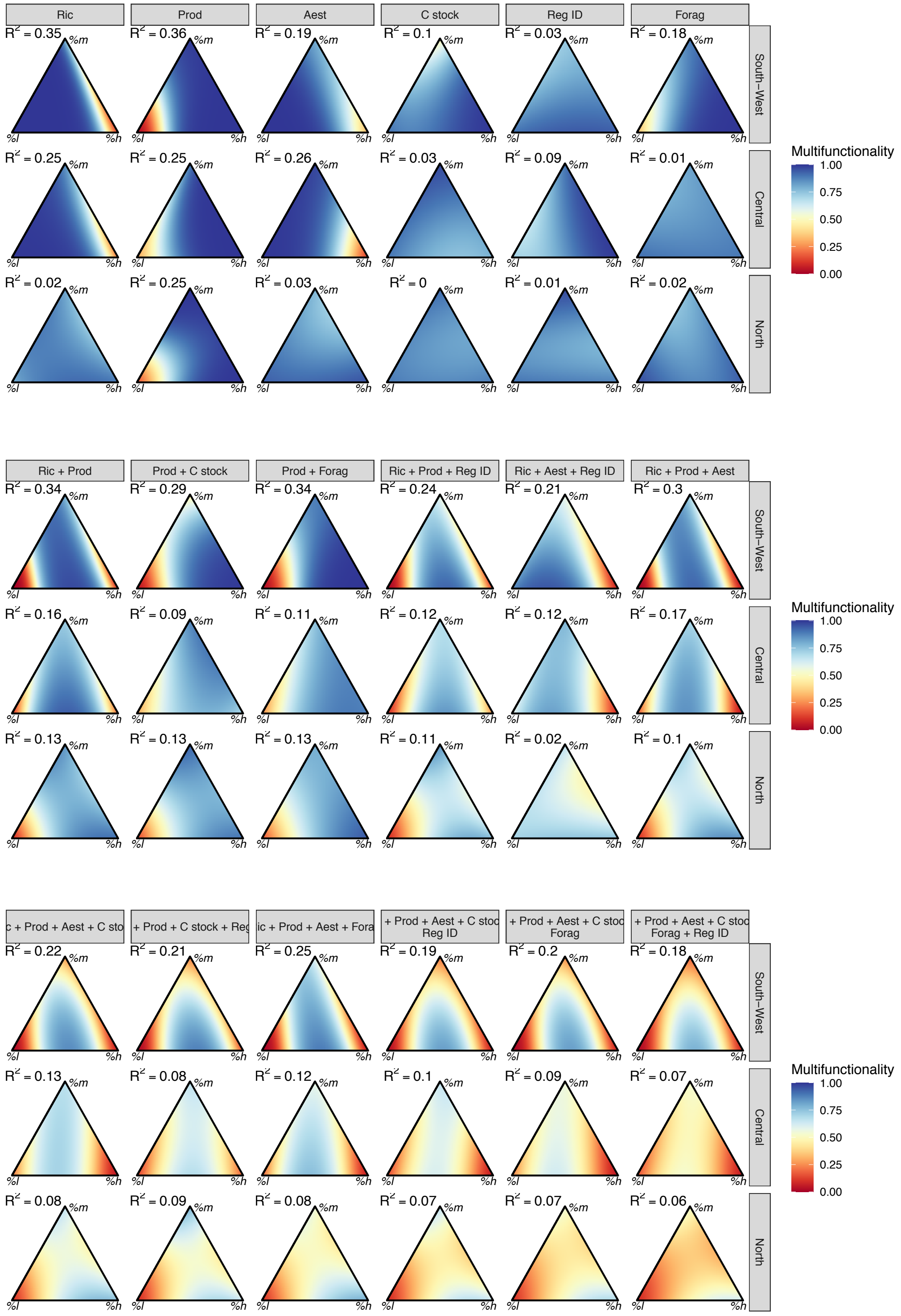

**Figure C14 Estimated multifunctionality values depending on landscape composition (in proportions of low, medium and high-intensity sites).** This figure differs from Fig. 5 in that the multifunctionality was calculated as 1 if all the services were above a 35th percentile threshold, and 0 otherwise (instead of a 25th percentile threshold).

The value presented corresponds to the probability that all given services are above the threshold. Blue indicates higher multifunctionality values, orange lower.

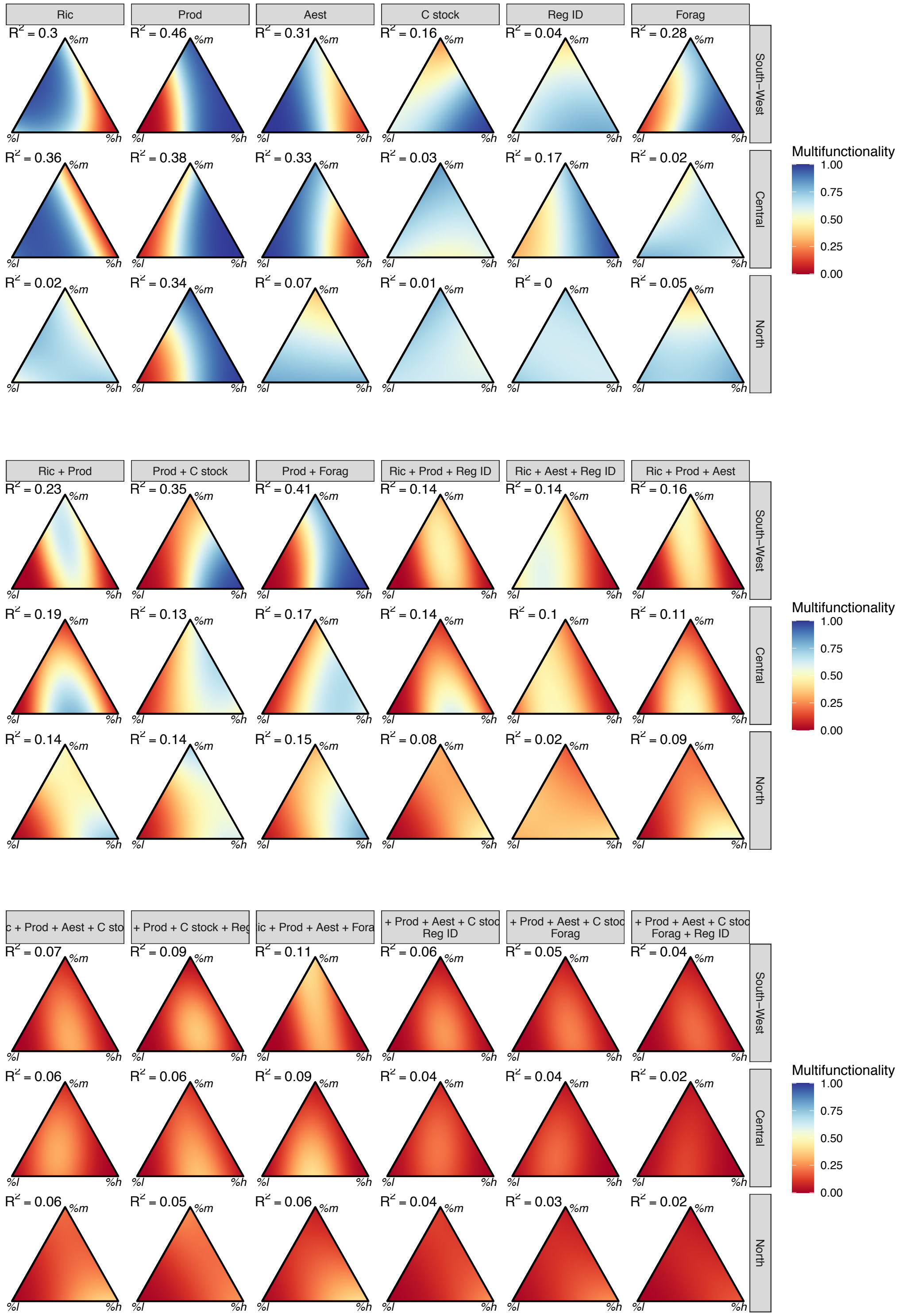

**Figure C15 Estimated multifunctionality values depending on landscape composition (in proportions of low, medium and high-intensity sites).** This figure differs from Fig. 4 in that landscapes multifunctionality was calculated as the average of the (scaled) values of all considered services, instead of the proportion of services above a threshold.

Blue indicates higher multifunctionality values, orange lower.

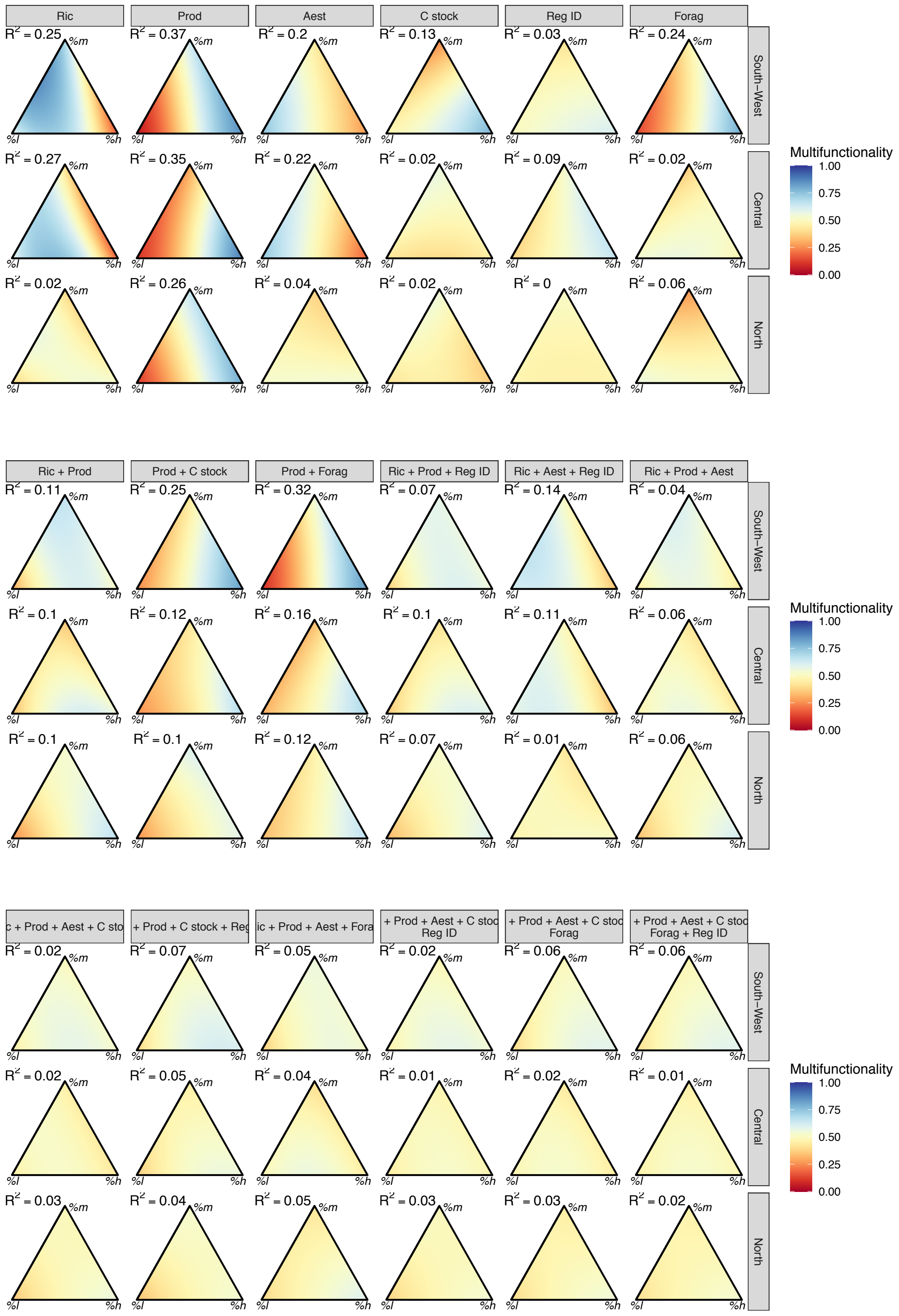

**Figure C16 Estimated multifunctionality values depending on the mean (x-axis) and coefficient of variation (y-axis) of the land-use intensity in the landscape**. The area outside the coloured represent combinations of intensity mean and variation that were not observed within the region.

Blue indicates higher multifunctionality values, orange lower.

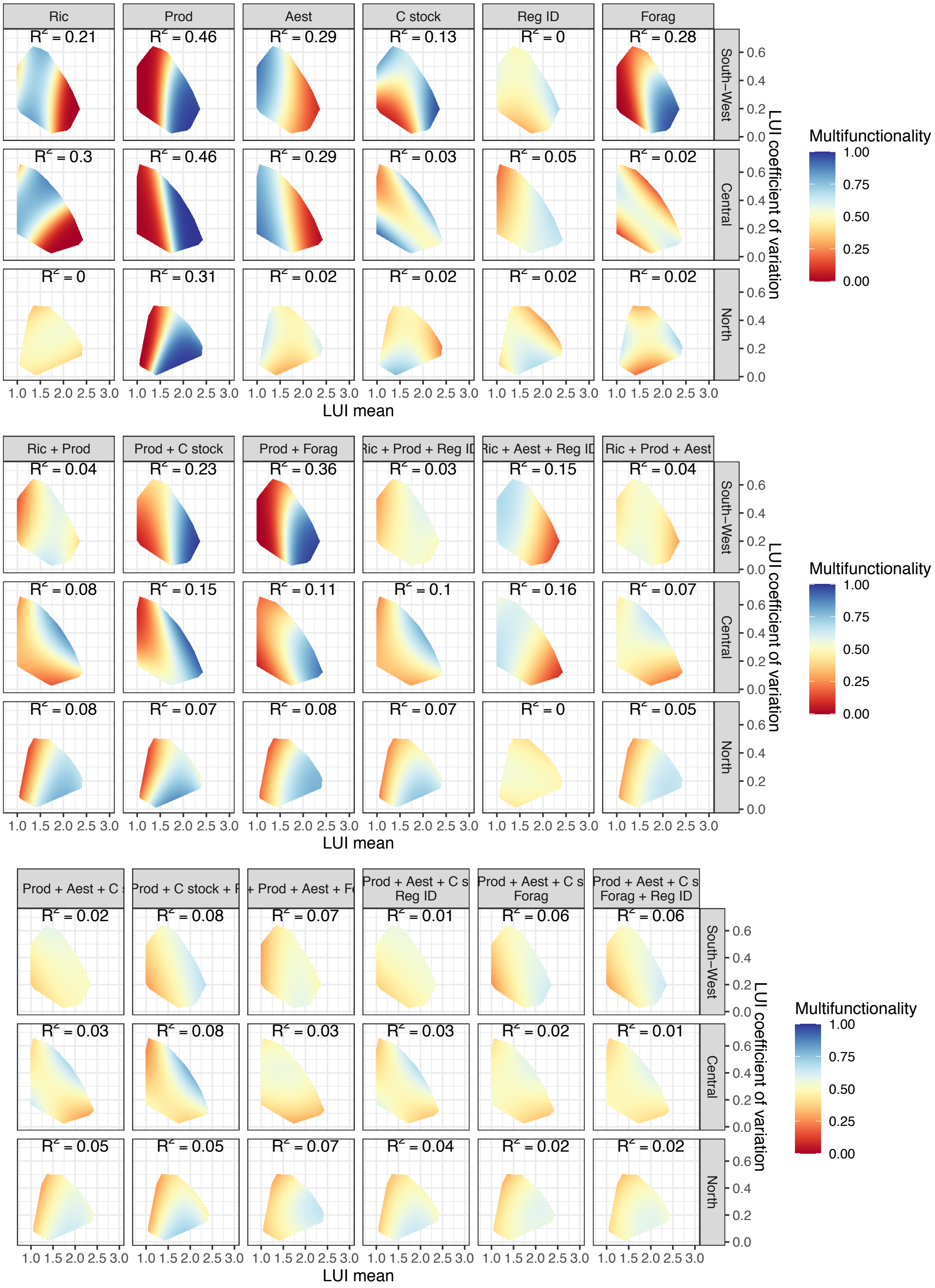

**Table C2** Variability of multifunctionality responsiveness with the ratio of the effect of intensity and other environmental variables ("Ratio"), the service response variance ("SRV"), and the number of services included in the analysis, depending on model parameters. Model results for ratio and number of services are presented as slope of the relationship ± standard deviation (* P> 0.05, ** P < 0.01, *** P< 0.001). For conciseness, model results for SRV include slope for the 'high SRM' group (service variance mean > median); the p-value is the significance of the interaction SRV*SRM. Sensitivity analyses include whether sites were classified into low-, medium- and high intensity s using all intensity values (quantile:30%) or only the lowest, middle and highest fifths (quantile: 20%); the number of sites per landscape (7, 10 or 13); the calculation of services at the landscape scale (either sum or max of the values observed at the site-level). Are also considered the different methodology to measure multifunctionality (proportion of services above a given threshold, either based on quantiles of the distribution of percentage of the maximum observed vales; average of service values; or as 1 if all services are above a threshold, 0 otherwise). Note that for the ratio of land use effect to other environmental variables, only models for service values not corrected for the environment are relevant and presented here.

| Env. correction | Intensity classes | N°of sites | Landscape-scale ES | Multifunctionality | Threshold | Ratio | SRV (high SRM) | N° of services |
|---|---|---|---|---|---|---|---|---|
| no | 20% quantile | 7 | mean | methodThreshold0.5 | 0.5 | 0,22+/-0,05*** | -0,85+/-0,1*** | -0,09+/-0,01*** |
| no | 30% quantile | 7 | mean | methodThreshold0.5 | 0.5 | 0,25+/-0,05*** | -0,79+/-0,09*** | -0,1+/-0,01*** |
| no | 30% quantile | 10 | max | methodThreshold0.5 | 0.5 | 0,13+/-0,06 | -0,49+/-0,08*** | -0,06+/-0,01*** |
| no | 30% quantile | 10 | mean | methodAverage | NA | 0,22+/-0,04*** | -0,53+/-0,07*** | -0,07+/-0,01*** |
| no | 30% quantile | 10 | mean | methodminimise | 0.15 | 0,28+/-0,05*** | 0,11+/-0,06*** | 0,01+/-0,01 |
| no | 30% quantile | 10 | mean | methodminimise | 0.25 | 0,23+/-0,05*** | -0,08+/-0,08*** | -0,06+/-0,01*** |
| no | 30% quantile | 10 | mean | methodminimise | 0.35 | 0,22+/-0,05*** | -0,31+/-0,11*** | -0,13+/-0,01*** |
| no | 30% quantile | 10 | mean | methodThreshold0.4 | 0.4 | 0,24+/-0,06*** | -0,64+/-0,09*** | -0,09+/-0,01*** |
| no | 30% quantile | 10 | mean | methodThreshold0.5 | 0.5 | 0,24+/-0,05*** | -0,81+/-0,1*** | -0,1+/-0,01*** |
| no | 30% quantile | 10 | mean | methodThreshold0.6 | 0.6 | 0,23+/-0,05*** | -0,7+/-0,09*** | -0,1+/-0,01*** |
| no | 30% quantile | 13 | mean | methodThreshold0.5 | 0.5 | 0,22+/-0,06** | -0,77+/-0,11*** | -0,11+/-0,01*** |
| yes | 20% quantile | 7 | mean | methodThreshold0.5 | 0.5 | NA | -0,53+/-0,11* | -0,08+/-0,01*** |
| yes | 30% quantile | 7 | mean | methodThreshold0.5 | 0.5 | NA | -0,4+/-0,09*** | -0,07+/-0,01*** |
| yes | 30% quantile | 10 | max | methodThreshold0.5 | 0.5 | NA | -0,15+/-0,09*** | -0,08+/-0,01*** |
| yes | 30% quantile | 10 | mean | methodAverageNA | NA | NA | -0,23+/-0,07*** | -0,05+/-0,01*** |
| yes | 30% quantile | 10 | mean | methodminimise | 0.15 | NA | 0,14+/-0,09*** | 0,02+/-0,01 |
| yes | 30% quantile | 10 | mean | methodminimise | 0.25 | NA | -0,05+/-0,09*** | -0,05+/-0,01*** |
| yes | 30% quantile | 10 | mean | methodminimise | 0.35 | NA | -0,22+/-0,12*** | -0,1+/-0,01*** |
| yes | 30% quantile | 10 | mean | methodThreshold0.4 | 0.4 | NA | -0,26+/-0,1*** | -0,08+/-0,01*** |
| yes | 30% quantile | 10 | mean | methodThreshold0.5 | 0.5 | NA | -0,29+/-0,1*** | -0,08+/-0,01*** |
| yes | 30% quantile | 10 | mean | methodThreshold0.6 | 0.6 | NA | -0,22+/-0,1*** | -0,07+/-0,01*** |
| yes | 30% quantile | 13 | mean | methodThreshold0.5 | 0.5 | NA | -0,29+/-0,11*** | -0,1+/-0,01*** |
